# Adaptation and correlated fitness responses over two time scales in *Drosophila suzukii* populations evolving in different environments

**DOI:** 10.1101/749945

**Authors:** Laure Olazcuaga, Julien Foucaud, Mathieu Gautier, Candice Deschamps, Anne Loiseau, Nicolas Leménager, Benoit Facon, Virginie Ravigné, Ruth A. Hufbauer, Arnaud Estoup, Nicolas O. Rode

**Affiliations:** 1CBGP, INRAE, CIRAD, IRD, Institut Agro, Université Montpellier, Montpellier, France; INRAE, UMR Peuplements Végétaux et Bio-agresseurs en Milieu Tropical, La Réunion, France; UMR PVBMT, CIRAD, St Pierre, La Réunion, France; Colorado State University, Department of Agricultural Biology and Graduate Degree Program in Ecology, Fort Collins, CO, USA

**Keywords:** Adaptation, experimental evolution, fitness landscape, fitness trade-off, specialization, reciprocal transplant experiment

## Abstract

The process of local adaptation involves differential changes in fitness over time across different environments. While experimental evolution studies have extensively tested for patterns of local adaptation at a single time point, there is relatively little research that examines fitness more than once during the time course of adaptation. We allowed replicate populations of the fruit pest *Drosophila suzukii* to evolve in one of eight different fruit media. After five generations, populations with the highest initial levels of maladaptation had mostly gone extinct, whereas experimental populations evolving on cherry, strawberry and cranberry media had survived. We measured the fitness of each surviving population in each of the three fruit media after five and after 26 generations of evolution. After five generations, adaptation to each medium was associated with increased fitness in the two other media. This was also true after 26 generations, except when populations that evolved on cranberry medium developed on cherry medium. These results suggest that, in the theoretical framework of a fitness landscape, the fitness optima of cherry and cranberry media are the furthest apart. Our results show that studying how fitness changes across several environments and across multiple generations provides insights into the dynamics of local adaptation that would not be evident if fitness were analyzed at a single point in time. By allowing a qualitative mapping of an experimental fitness landscape, our approach will improve our understanding of the ecological factors that drive the evolution of local adaptation in *D. suzukii*.

## Introduction

Adaptation to local environmental conditions plays a central role in the maintenance of biodiversity (Levins 1968; Felsenstein 1976; Gillespie and Turelli 1989). In particular, when selection is divergent across environments, if migration is low enough, rare genotypes can accumulate in environments where they have the highest fitness (Deakin 1966; Gillespie 1975; Svardal et al. 2015). Despite the importance of understanding this process of local adaptation, its investigation in natural populations has been limited by a number of experimental and conceptual factors (Rausher 1988; Fry 1996; Hansen et al. 2006; Barghi et al. 2020). In particular, disentangling natural selection from genetic drift requires accurate fitness estimates that are relevant to natural environments, a notoriously difficult task (Fry 1996; Forister et al. 2012). Heterogeneity in environmental conditions and lack of knowledge regarding the starting populations represent additional challenges to studying the process of adaptation in natural populations (Barghi et al. 2020).

Experimental evolution under laboratory conditions represents a powerful alternative to studying the dynamics of local adaptation in the field (Fry 2003; Kawecki et al. 2012). The approach consists of allowing replicated experimental populations to evolve in different environments and performing reciprocal transplant experiments to study fitness changes in each environment. Here, we call the environment in which a population is evolving the selective environment. Reciprocal transplant experiments following experimental evolution can reveal whether fitness changes in the selective environment (i.e., “direct fitness reponses” sensu Bennet et al 1992) are greater than fitness changes in other environments (i.e., “correlated fitness responses” sensu Bennet et al 1992). These experiments also can be used to test directly for a pattern of local adaptation (Blanquart et al 2013) by investigating whether populations have higher mean fitness in the selective environment (in this context also often called the sympatric environment) than in other environments (the non-selective or allopatric environments). To emphasize that selective environments determine the selective pressures at the origin of the process of local adaptation, we will hereafter solely use the terms selective and alternative environments.

Evidence for the evolution of local adaptation has been mixed in experimental evolution studies over a single time scale (for reviews see Fry 1996; Kassen 2002; Hereford 2009; Jasmin and Zeyl 2013; Bono et al. 2017; Bergh et al. 2018). For example, adaptation to one environment can be associated with an increase (e.g., Bennett et al. 1990; Magalhães et al. 2009; Messina et al. 2009; Laukkanen et al. 2012; Messina and Durham 2013), a decrease (e.g., Fry 1990; Mackenzie 1996; Turner and Elena 2000; Agudelo-Romero et al. 2008; Bedhomme et al. 2012; Messina and Durham 2015; Gompert and Messina 2016) or no change (e.g., Fry 2001) in fitness in other environments. These contrasting results are not actually contradictory if interpreted in the light of evolutionary theory, particularly that based on invasion fitness and evolutionary branching (Svardal et al. 2015) or based on fitness landscapes theory and Fisher’s geometric model (Martin and Lenormand 2015). If a population has split into subpopulations that experience two different environments with low migration, whether or not adaptation to the selective environment leads to an increase in fitness in the alternative environment will depend upon how similar those environments are to each other and to the ancestral environment (Fig. 1). Specifically, whether adaptation to the selective environment leads to adaptation to other environments is highly dependent on i) the (mal)adaptation of the ancestral population to the two selective environments and ii) the amount of time a population has spent in the selective environment. If the phenotypes that maximize fitness in the two new environments are similar to each other, but relatively different from the optimal phenotype in the ancestral environment, adaptation will increase in both environments regardless of how long a population has experienced an environment (Fig. 1A). In contrast, if the optimal phenotypes in the two new environments are different from each other, but more similar to each other than they are to the optimal phenotype in the ancestral environment, whether or not adaptation to the selective environment leads to an increase in fitness in the alternative environment will depend on when fitness is measured because selection in the two environments changes over time from being convergent to divergent (Fig. 1B). During the first few generations, adaptive evolution will shift the mean phenotype in each environment in a similar direction, so that adaptation increases in both environments (step 1 in Fig. 1B); but over time, those mean phenotypes will diverge, resulting in adaptation in one environment and maladaptation in the other environment (step 2 in Fig. 1B). This scenario shown in Fig. 1B is often not explicit in theoretical models (e.g., Levins 1962), which often focus on predicting the outcome of evolution rather than the dynamics of evolution (Roughgarden 1979). Consistent with the scenario shown in Fig. 1B, a temporal reversal in the association between adaptation in a given selective environment and fitness changes in alternative environments has been observed in experimental populations of yeast (Jasmin and Zeyl 2013) and bacteria (Satterwhite and Cooper 2015; Schick et al. 2015). Also consistent with Fig. 1B is the observation that negative associations between the rate of adaptation in an environment and the rate of fitness decline in other environments are more likely to be found in experimental evolution studies performed over about 10,000 generations than in studies performed over fewer generations (Bono et al. 2017).

**Figure 1.**
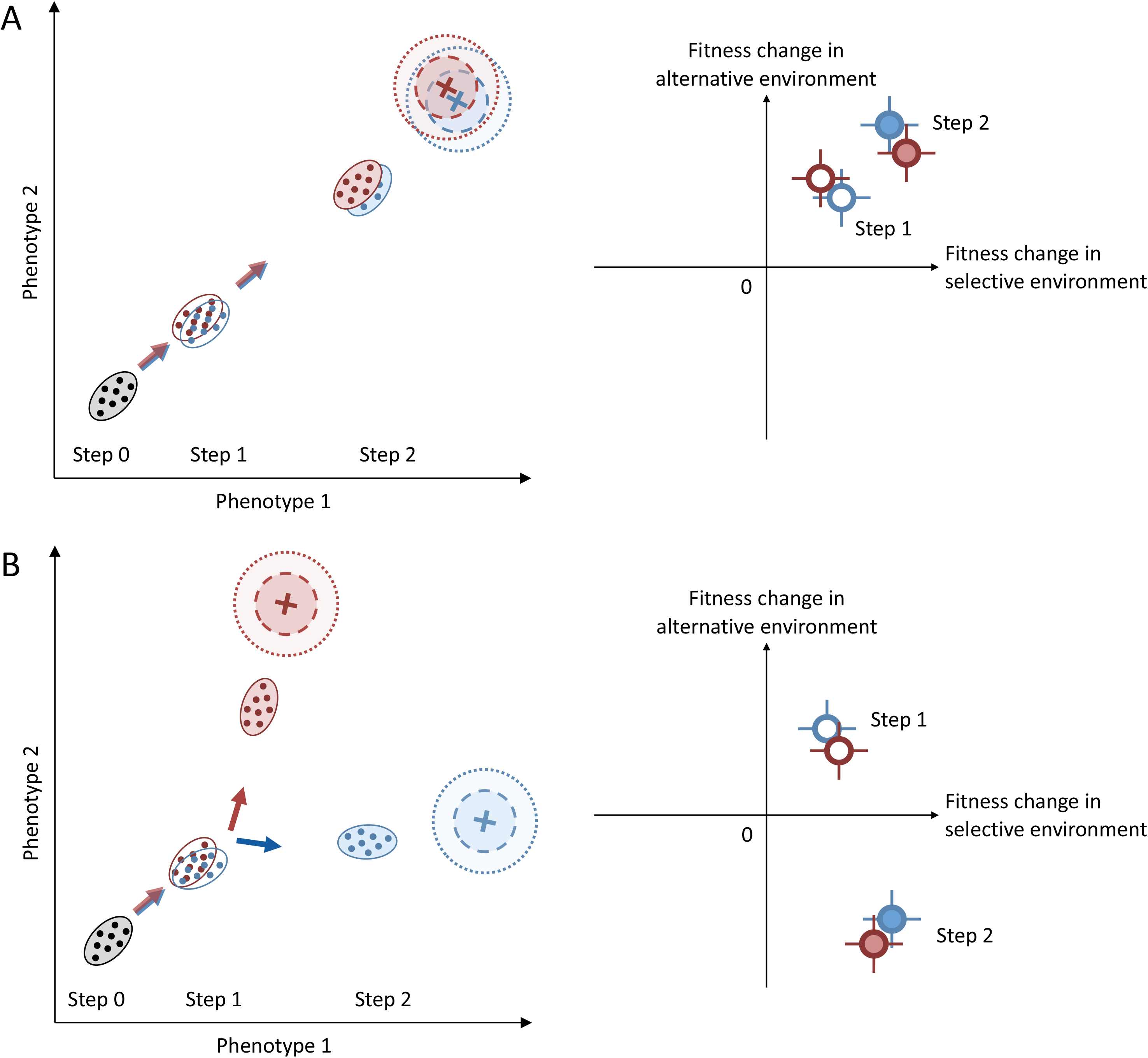
Changes in fitness in selective and alternative environments depend on the level of adaptation of the ancestral population and on the relative positions of the phenotypic optima. In a two-dimensional fitness landscape, phenotypic optima for environments 1 and 2 (red and blue crosses respectively) are either (A) closed to each other or (B) distant from each other. In the left panels, the ancestral population, the population evolving in environments 1 and the population evolving in environments 2 are represented by grey, red and blue ellipses, respectively. In the right panels, the fitness changes relative to the ancestral population (step 0) in the selective environment or in the alternative environment are represented for step 1 and 2, respectively. Open and closed ellipses respectively correspond to early (step 1) and late (step 2) adaptation.

In all, we found only a handful of studies that evaluated adaptation and correlated responses over more than one time period to test whether selection patterns change from convergent to divergent (spider mite: Magalhães et al. 2009; yeast: Jasmin and Zeyl 2013; bacteria: Satterwhite and Cooper 2015; Schick et al. 2015; virus: Agudelo-Romero et al. 2008). Most of them considered adaptation to different environments of microorganisms grown from a single clone to different environments (Jasmin and Zeyl 2013; Satterwhite and Cooper 2015; Schick et al. 2015). However, many natural populations of interest are macro-organisms, in which population sizes are smaller and generation times longer than for experimental micro-organisms. In such populations, standing genetic variation can be expected to play a larger role in adaptation to a new environment than *de novo* mutations (Barrett and Schluter 2008, Bailey & Bataillon 2016). Although providing an important jumping off point, microbial studies provide an incomplete picture of the process of local adaptation from standing genetic variation. The spider mite study addressed this issue using genetically diverse spider mite populations, and found a positive association between adaptation to a given environment and fitness changes in other environments after both 15 and 25 generations (Magalhães et al. 2009). This result suggests that populations adapted to two new environments whose fitness optima were close to each other, as represented in Fig. 1A. Studies that investigate adaptation and correlated responses to selection in genetically diverse populations adapting to a wide variety of environments such that some are likely to be different from each other and from the ancestral environment (Fig. 1B) are deeply needed.

The spotted wing drosophila, *Drosophila suzukii* Matsumura (Diptera: Drosophilidae), represents an attractive biological model to address this issue. First, growth media made of different fruits can be used to represent environments with different fitness optima (Olazcuaga et al. 2019). When comparing natural *D. suzukii* populations sampled in different fruits, Olazcuaga (2019) recently found that emergence rates were significantly higher on fruit media corresponding to the fruit from which each population originated, than on fruit media corresponding to alternative fruits. This suggests that the standing genetic variation that segregates in natural populations is sufficient to adapt to different fruit media during experimental evolution in the laboratory. In addition, *D. suzukii* presents several advantages for experimental evolution including short generation time, small size, and relative ease of maintenance of large populations over many generations in the laboratory. These characteristics greatly facilitate a straightforward estimation of population mean fitness across different environments, using the same protocol as the one used for experimental evolution.

Consequently, this biological system does not rely on phenotypic traits that might be only loosely associated with fitness in experimentally evolving populations. Finally, a better understanding of the potential of *D. suzukii* to adapt to the fruits of different host plants is of agronomic interest, as this species is a major pest of berries and stone fruits (Asplen et al. 2015). In this study, we maintained *D. suzukii* populations on different fruit media and investigated adaptation to those different media as well as fitness changes on alternative media using reciprocal transplant experiments after five and 26 generations of evolution. We specifically addressed the following questions: 1) Are populations able to adapt to all selective environments at the same rate (i.e., are direct fitness responses positive and of the same magnitude across environments)? 2) Are fitness changes in selective and alternative environments of the same sign and magnitude across replicate populations evolving on the same fruit medium? 3) Is there a reversal over time in the direction of the association between fitness changes in selective and alternative environments?

## Materials and methods

### Field sampling

To initiate a laboratory population, approximately 1,000 *D. suzukii* flies (more than 500 females and 460 males) were collected using baited cup traps (Lee et al. 2013) at six sampling sites within 10 km of Montpellier, France in September and October 2016. At this time of the year, most of the fruit crops cultivated over large areas (e.g., cherries and strawberries) are no longer available. Based on the literature (Poyet et al. 2015), we estimate that flies have potentially emerged from more than 18 wild (i.e., non crop) host plants from nine families available in the sampling area in October (Table S1). Thus our sampled individuals likely emerged from a large range of host plants rather than a few specific host plants, enabling us to establish a laboratory population likely to harbor alleles that could potentially provide adaptation to a large variety of host plants.

### Laboratory maintenance

As we did not want to create a population adapted to a particular fruit, flies were maintained for nine generations in 10 ml vials with standard laboratory fly food (consisting of sugar, dry yeast, minerals and antifungal solution; Backhaus et al. 1984), prior to the start of the experiment. This maintenance on a protein-rich medium likely selected for a combination of phenotypic traits different from those selected on host fruits (which are likely to be poor in proteins). Each generation, newly emerged adults were mixed across vials and randomly distributed into groups of 20 adults in 100 new vials to maintain a large panmictic population (i.e., ∼2000 individuals) and as much genetic variation as possible. We chose this approach rather than using isofemale lines to maintain genetic variation because linkage disequilibrium in synthetic populations recomposed by crossing isofemale lines decreases additive genetic variance (Kessner and Novembre 2015) and the subsequent response to selection. In addition, selection among isofemales due to different degrees of inbreeding depression could reduce genetic diversity at loci responsible for adaptation to different fruits. Our large population of flies was maintained with discrete generations over two-week cycles (21°C, 65%, 16:8 day/night light cycle; see Olazcuaga et al. 2019 for details). Experimental evolution and phenotyping were performed under the same temperature, humidity and light conditions.

### Fruit media

To investigate the dynamics of fitness changes in selective and alternative environments, we reared replicate populations on media made using fruit purees. To standardize generation time and fitness estimation protocol across environments, we compare emergence rates across fruit media (Fig. 4 in Olazcuaga et al. 2019). We chose eight fruit media where our population exhibited similar emergence rates (blackcurrant, cherry, cranberry, fig, grape, strawberry, rose hips, tomato), and which correspond to host fruits attacked to varying degrees in the field (Walsh et al. 2011; Cini et al. 2012; Bellamy et al. 2013; Steffan et al. 2013; Kenis et al. 2016; Kanda et al. 2019).

To limit variation within each environment throughout the experiment and to directly compare fruits that ripen at different times of the year, we used media made with frozen fruit purees rather than whole fruits (recipe available in Olazcuaga et al. 2019). We hence assume that temporal variation within each environment is minimal so that we can compare fitness across phenotyping steps. We also assume that fruit media differ in their biochemical properties and select for different phenotypic optima. This assumption seems reasonable, as adaptation to some of these media varies across natural populations and matches the fruit from which they originated (Olazcuaga 2019).

### Experimental evolution experiment

The three phases of the evolution experiment are summarized in Fig. 2. During phase 1, we established experimental populations by deriving replicate populations from our base population and placing them on each of eight different fruit media (Fig. 2). Five replicate populations were established per fruit (40 populations in total). Each population consisted of 400 adults (20 vials of 20 flies), which corresponds to a reasonably large population size to limit genetic drift (Woodworth et al. 2002). Populations were maintained on a 21-day cycle. For each vial, we placed 20 six-day-old flies into a vial filled with 10 ml of a single fruit medium to mate and oviposit. At this stage of adult development, all adult females should be ready to oviposit (Emiljanowicz et al. 2014). The sex ratio of the adults was neither controlled nor measured. While not controlling sex ratio increased variation in the total number of eggs among tubes, it made it possible to handle more flies, increase replication, and minimize additional stress on the flies due to manipulation (for instance a longer time of CO_2_ anesthesia). The sex ratio did not evolve over the course of the experiment (Olazcuaga 2019). After 18 hours, adults were removed. After 15 days, we anesthetized emerging adults from each replicate population using CO2, mixed them across vials to produce 20 groups of 20 flies, each placed in a vial with 5 ml of the same fruit medium. Adults could mature for 6 days before starting the next cycle. At most 400 individuals per population were kept to produce the next generation. If fewer than 400 individuals emerged, less than 20 vials were made but always with 20 adults each. Populations were randomly distributed among eight racks (100 vials per rack, composed of five populations of 20 vials each) and randomly arranged spatially in a climate chamber. The 40 populations were reared in two temporal blocks separated by two days.

**Figure 2.**
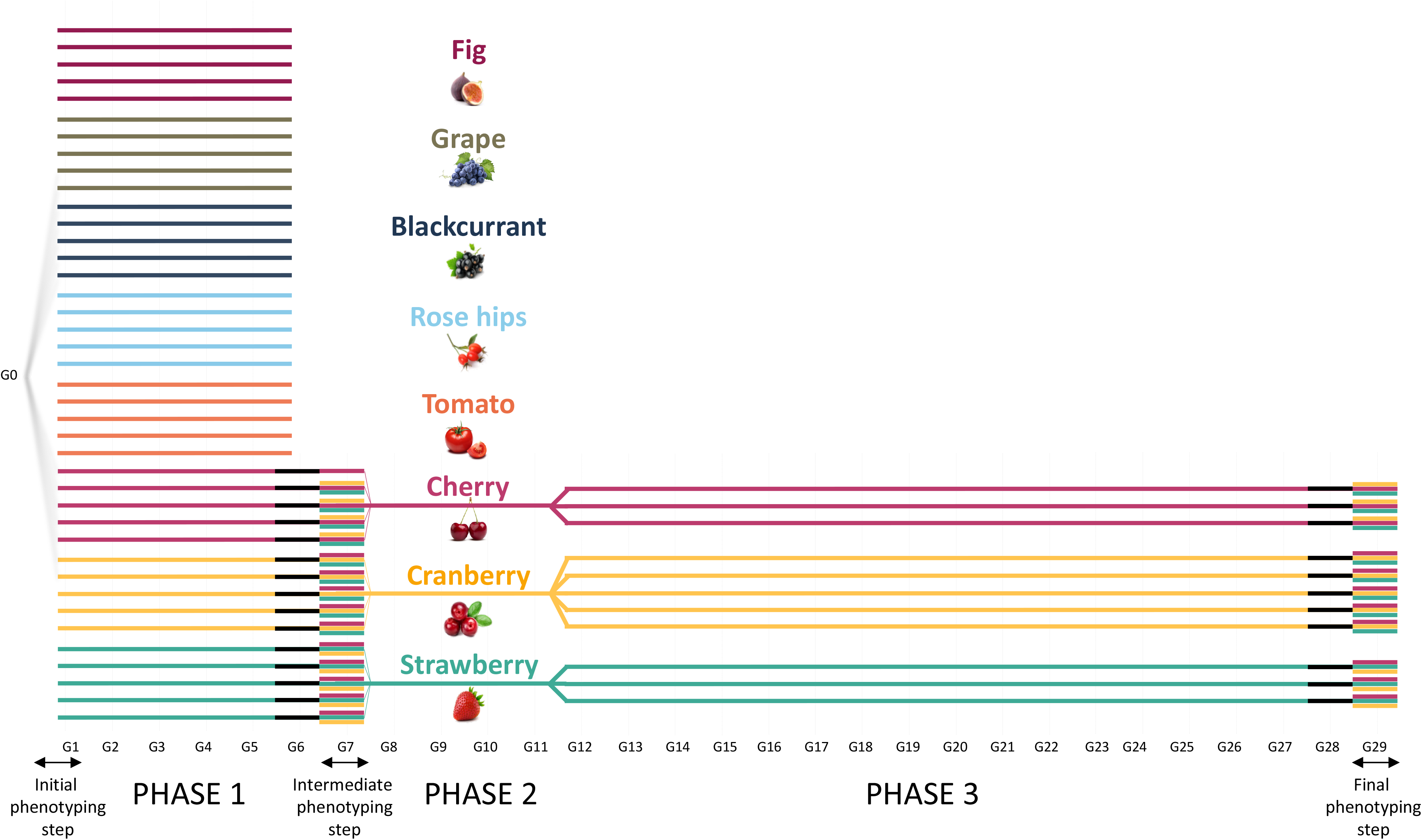
Experimental evolution design depicting the different fruit media and the three phenotyping steps. Before each phenotyping step, populations spend one generation in a common environment (standard laboratory fly medium), represented by a black line. For each phenotyping step and each population, we measured fitness in each of the three fruit media with an average of 9 (range: 2-32) and 30 vials for the intermediate and final phenotyping steps, respectively. During the intermediate phenotyping step, fitness on alternative environment was not measured for one replicate population evolving on cherry due to small population size.

By the fifth generation of experimental evolution, populations on blackcurrant, fig, grape, rose hips or tomato were either extinct or close to extinction, with fewer than 30 individuals per population (4 to 28 individuals; see Appendix S1, Fig. S1). Experimental populations persisted over the first five generations on cherry, cranberry and strawberry, but some of the replicates had relatively small population sizes (mean of the population size after five generations on fruit ± sd for populations on cherry, cranberry and strawberry: 85.0 ± 62.0, ± 58.5 and 133.8 ± 134.8, respectively). During phase 2 (seven generations after the start of the experiment), the five replicate populations on each fruit were pooled together to counteract inbreeding depression that we assumed to be the main driver of the decline in population size (1840, 420 and 2220 individuals pooled for cranberry, cherry and strawberry respectively, with demographic and genetic stochasticity likely being responsible for differences across fruit media).

During phase 3 (11 generations after the start of the experiment), the three pooled populations had recovered (3040, 1560 and 1600 individuals for cranberry, cherry and strawberry respectively) and were divided into five, three and three replicate populations for cranberry, cherry and strawberry, respectively (with 500 individuals per replicate population; Fig. 2). The number of replicate populations per fruit depended on the number of individuals available. During this third phase of experimental evolution, each population was maintained at a size of 500 individuals (25 tubes of 20 individuals) using the same protocol as described above for the first phase. Populations were randomly distributed among four racks (75 vials per rack i.e., three populations of 25 vials). The 11 populations were reared in a single temporal block. To estimate Malthusian fitness at each generation, we counted the number of adults that emerged from each vial (except during the pooling step where we counted the total number of adults across all vials and computed an average fitness across vials).

To reduce experimental burden, we compared the fitness of evolved populations with that of the ancestral population rather than to that of an evolved control population. As fruit media were based on the same stock of frozen puree, we assume the variation in experimental conditions across phenotyping steps to be small. Moreover, experiments using inbred lines show that variation among replicates within a single phenotyping assay is as large as variation between phenotyping assays several generations apart (Olazcuaga 2019). In addition, because the combination of host fruits used by the population sampled in the wild was unknown, no single fruit medium could represent an appropriate control for our experiment. Finally, using a control population maintained on standard medium (as recommended by Fry 2003) was likely inappropriate. Our ancestral population was likely adapted to our rearing schedule (discrete generations over three weeks) but might not have been entirely adapted to the standard medium. As a result, control populations maintained on standard medium would have likely diverged from the ancestral population over the course of the experiment. Small populations would have experienced genetic drift while large populations would have experienced selection.

### Estimation of fitness changes in selective and alternative environments using reciprocal transplant experiments

We estimated the average fitness of each population in each of the three fruit media (cherry, cranberry and strawberry) during the initial, intermediate and final phenotyping steps (respectively corresponding to one, seven and 29 generations after the start of the experiment; Fig. 2), after one generation in a common garden (standard laboratory medium to standardize maternal environmental effects; Fry 2003). Thus, populations had evolved for five and 26 generations in each selective fruit medium when their fitness was estimated during the intermediate and final phenotyping steps respectively. In phenotyping we used the same protocol as that used to maintain experimental populations and estimated fitness over the 21- day life cycle (number of adults emerging that descended from the 20 initial adults from the previous generation). We also counted the number of eggs in vial so that our fitness measure corresponded to the product of number of eggs and egg-to-adult viability. During the initial phenotyping step, the average fitness of the base population in each fruit medium was estimated using 100 vials (two temporal blocks). The average fitness of each evolved population was estimated in each fruit medium using between two and 32 vials during the intermediate phenotyping step and 30 vials during the final phenotyping step (three and six temporal blocks, respectively).

### Statistical analyses

All analyses were performed using the R statistical software (R Core Team 2014).

#### Fitness change in selective environments

To investigate the temporal dynamics of adaptation, we used census data recorded each generation during the three phases of the experiment. To avoid environmental effects (including maternal effects), we excluded data from generations where individuals or their parents developed in standard medium. As we used discrete generations, we computed the average Malthusian fitness of each population at the the *i*th generation as: 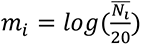, where *N*_i_ is the average number of emerging adults per tube over one life cycle. For each generation and each population, we considered the average fitness across vials and used the number of vials counted as a weight in the analyses. To test for differences in the rate of adaptation among fruit media, we fitted the following linear model on average fitness mijkl:

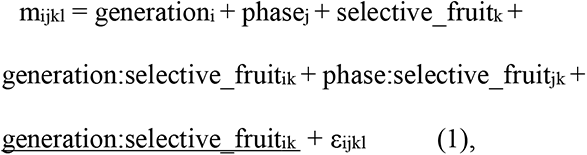

where fixed effects included the effect of the *i*th generation as a covariate (*generationi*, with *i*=2,..,27), the effect of the *j*th phase of experimental evolution (*phasej*, with either *j*=1,…,3), the effect of the *k*th selective fruit (*selective*_*fruitk*, with *i*=1,..,3 for cherry, cranberry and strawberry respectively), the interaction between the *generation* and *selective*_*fruit* effects, the interaction between the *phase* and *selective_fruit* effects. Random effects included the interaction between the *i*th generation and the *k*th *selective*_*fruit* (mean of zero and varianceσ^2^_gen fruit_) to control for potential batch effects among vials of the same fruit medium cooked across generations. A random error (ε_ijkl_ mean of zero and variance σ^2^_res_) accounted for the variation among populations evolving on the same fruit medium. We compared several simpler models derived from this full model that included none, one or more of the effects described above), while keeping random effects in all models. All the models tested are listed in Table 1. Due to their partial redundancy, the *generation* and *phase* effects were never tested together. Models were ranked according to their corrected Akaike’s information criterion (AICc; Hurvich and Tsai 1989) using the *MuMIn* package (Barton 2009). For each model, we computed ΔAICc value as the difference between the AICc of that model and the best fit model. Best competing models with AICc differences lower than two were considered as strongly supported by the data, except when they differed by a single degree of freedom and had essentially the same log-likelihood (Burnham and Anderson 2002).

**Table 1:**
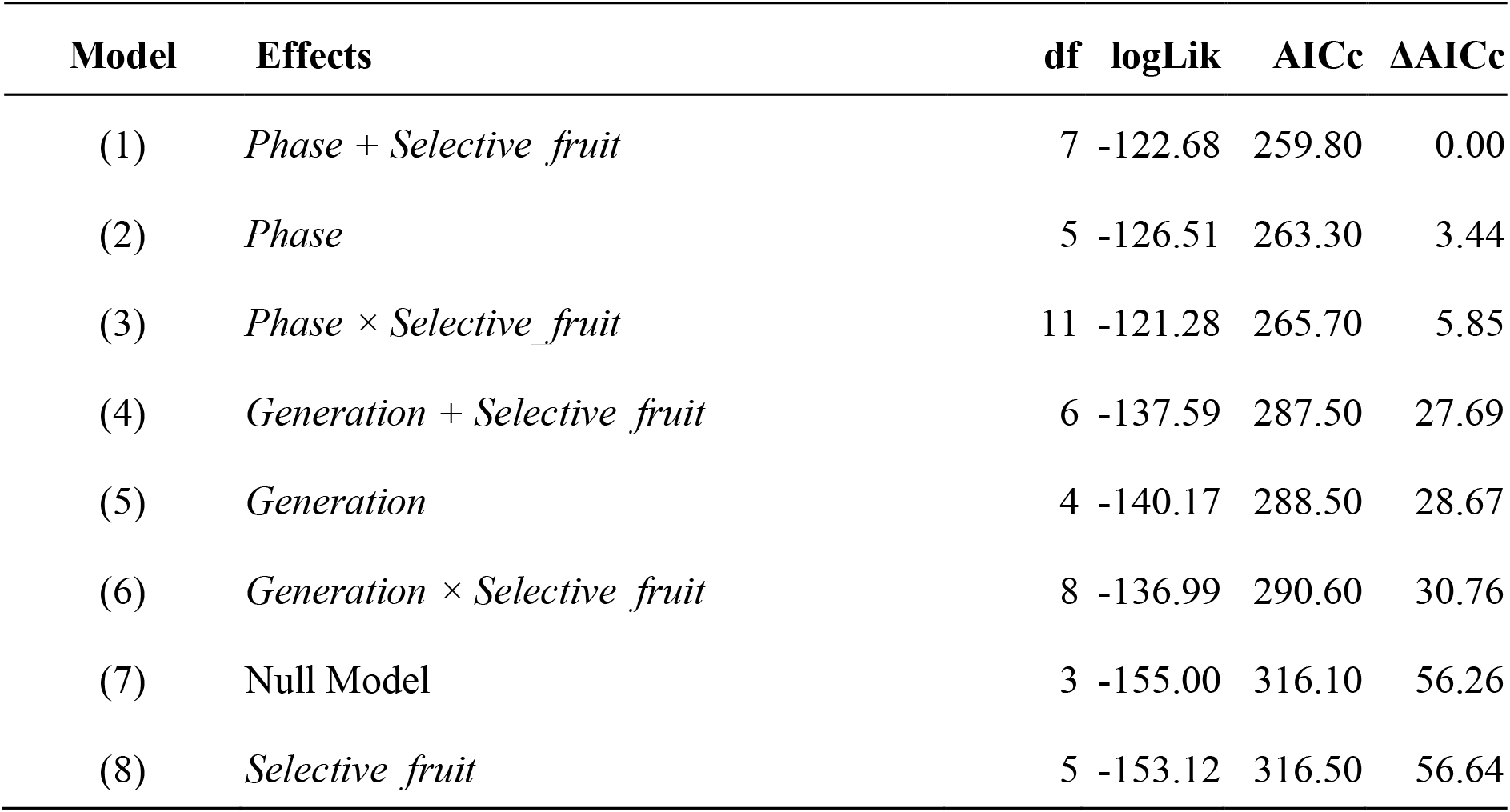
Results of the AICc model selection for comparing temporal dynamics of adaptation across the three selective fruits. All models included *generation:selective_fruit* as a random effect.

| Model | Effects | df | logLik | AICc | ΔAICc |
| --- | --- | --- | --- | --- | --- |
| (1) | *Phase + Selective_fruit* | 7 | -122.68 | 259.80 | 0.00 |
| (2) | *Phase* | 5 | -126.51 | 263.30 | 3.44 |
| (3) | *Phase × Selective_fruit* | 11 | -121.28 | 265.70 | 5.85 |
| (4) | *Generation + Selective_fruit* | 6 | -137.59 | 287.50 | 27.69 |
| (5) | *Generation* | 4 | -140.17 | 288.50 | 28.67 |
| (6) | *Generation × Selective_fruit* | 8 | -136.99 | 290.60 | 30.76 |
| (7) | Null Model | 3 | -155.00 | 316.10 | 56.26 |
| (8) | *Selective_fruit* | 5 | -153.12 | 316.50 | 56.64 |

Finally, a complementary analysis was performed on phases 1 and 3 separately to test whether the rate of adaptation differed across replicate populations evolving on the same fruit medium (see Appendix S1).

#### Comparison across populations of the direction of fitness changes in selective and alternative environments

To quantify the direction and magnitude of fitness changes in selective and alternative fruit media, we combined the data of the three phenotyping steps and fitted the following negative binomial model (log link) on the number of adults, nijkl, that emerged from each tube:

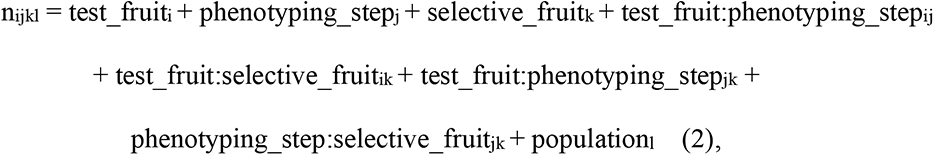

where fixed effects included the effect of the *i*th test fruit medium (*test_fruiti*, with i=1, 2 and 3, for cherry, cranberry and strawberry respectively), the effect of the *j*th phenotyping step (*phenotyping_step*, with either j=1, 2 and 3 for the initial, intermediate and final phenotyping steps, respectively), the effect of the *k*th selective fruit medium (*selective_fruitk*, with k=1, 2 and 3, for cherry, cranberry and strawberry respectively), and their two-way or and three-way interactions and a random population effect (population with mean of zero and variance σ^2^). We used the *glmer.nb* function of the *lme4* package and computed the 95% confidence interval of each parameter with 1,000 simulations using the *bootMer* function (Bates et al. 2015). Similarly, we used a negative binomial model and a binomial model to respectively investigate changes in fecundity (corresponding to count data) and in egg-to-adult viability (corresponding to proportion data) between phenotyping steps.

#### Correlation between fitness changes in selective and alternative environments

To investigate whether fitness changes in selective environments were associated with positive or negative fitness changes in alternative environments, we used two approaches: i) for populations from each pair of environments, we tested whether fitness changes in selective and alternative environments were in the same or in different directions and ii) we tested whether fitness changes in selective fruit media were significantly greater than fitness changes in alternative fruit media. First, for each pair of environments and for the intermediate and final phenotyping steps separately, we estimated the correlation coefficient between the average fitness change of populations in their selective medium and their average fitness change in each of the two alternative media using the sum of number of tubes of selective and fruit media as weight and estimated its 95% confidence interval using the *sjstats* package (Lüdecke 2018). For each pair of fruit media, we estimated the difference between the correlation coefficients estimated during the intermediate and final phenotyping steps and tested whether the 95% confidence interval of this difference included zero (Zou 2007). To illustrate the relationship between fitness change in selective and alternative fruit media, we estimated the intercept and the slope of the regression of fitness change in alternative environments over fitness change in selective environments using a Major Axis regression in the *lmodel2* package (Legendre 2014). This method requires the existence of variation across replicate populations and assumes that the correlation between fitness changes in selective and alternative environments is independent of the selective environment.

Second, to test whether fitness changes in selective fruit media were significantly greater than fitness changes in alternative fruit media, we used ‘sympatric–allopatric’ (SA) contrasts (Blanquart et al. 2013), a test that controls for variation among populations (e.g., due to different levels of inbreeding) and among environments (e.g., due to differences in environment quality). To this end, we fitted the following linear model on the average fitness change relative to the ancestor, s_ijk_, for the intermediate and final phenotyping steps separately:

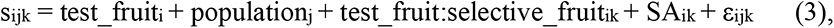

where fixed effects included the effect of the *i*th test fruit medium (*test_fruiti*, with i=1,…,3, for cherry, cranberry and strawberry respectively), the effect of the *j*th population (*populationj*, with either j=1,…,14 or j=1,…,11 for the intermediate and final phenotyping steps, respectively), the interaction between the *i*th test fruit and the *k*th selective fruit medium on which the *j*th population evolved (*selective_fruitk*, with k=1,…,3, for cherry, cranberry and strawberry respectively), a sympatric vs. allopatric effect that measures local adaptation (*SAik*) and a random error (ε_ijkl_ mean of zero and variance σ^2^_res_). We did not include a *selective_fruit* effect, which is already statistically accounted for by the *population* fixed effect. For each population, we used the inverse of the variance of each estimate (se(s_ijk_)^2^) as a weight in the analyses. To test for a pattern of local adaptation, we used a two-way ANOVA and computed a F-test with the appropriate degrees of freedom (eq. D1 in the supplementary information of Blanquart et al. 2013). We performed a set of computer simulations showing that, at least in our experimental setup (high replicate level and intermediate overdispersion of the count data), the *F*-test proposed by Blanquart et al. (2013) can be applied to count data (see Appendix S3).

## Results

### Temporal fitness change in each selective fruit medium

Fitness change in each selective fruit medium is shown for the three phases of the experiment in Fig. 3 and Fig. S1. During phase 1, fitnesses were initially negative in all eight fruit media and differed significantly across fruit media (ΔAICc > 28.3 for models 14 and 15 without a fruit effect, Tables S2), but not across populations evolving on the same fruit medium (ΔAICc = 9.48 for the model 12 including a population effect, Table S2). Temporal changes in fitness differed across fruits (ΔAICc = 2.12 for the model 10 without an interaction between the fruit and generation effects, Table S2). Populations with the lowest initial fitness (Table S3) went extinct by the fifth generation due to demographic effects, so that subsequent fitness changes could only be measured on cherry, strawberry and cranberry media.

**Figure 3.**
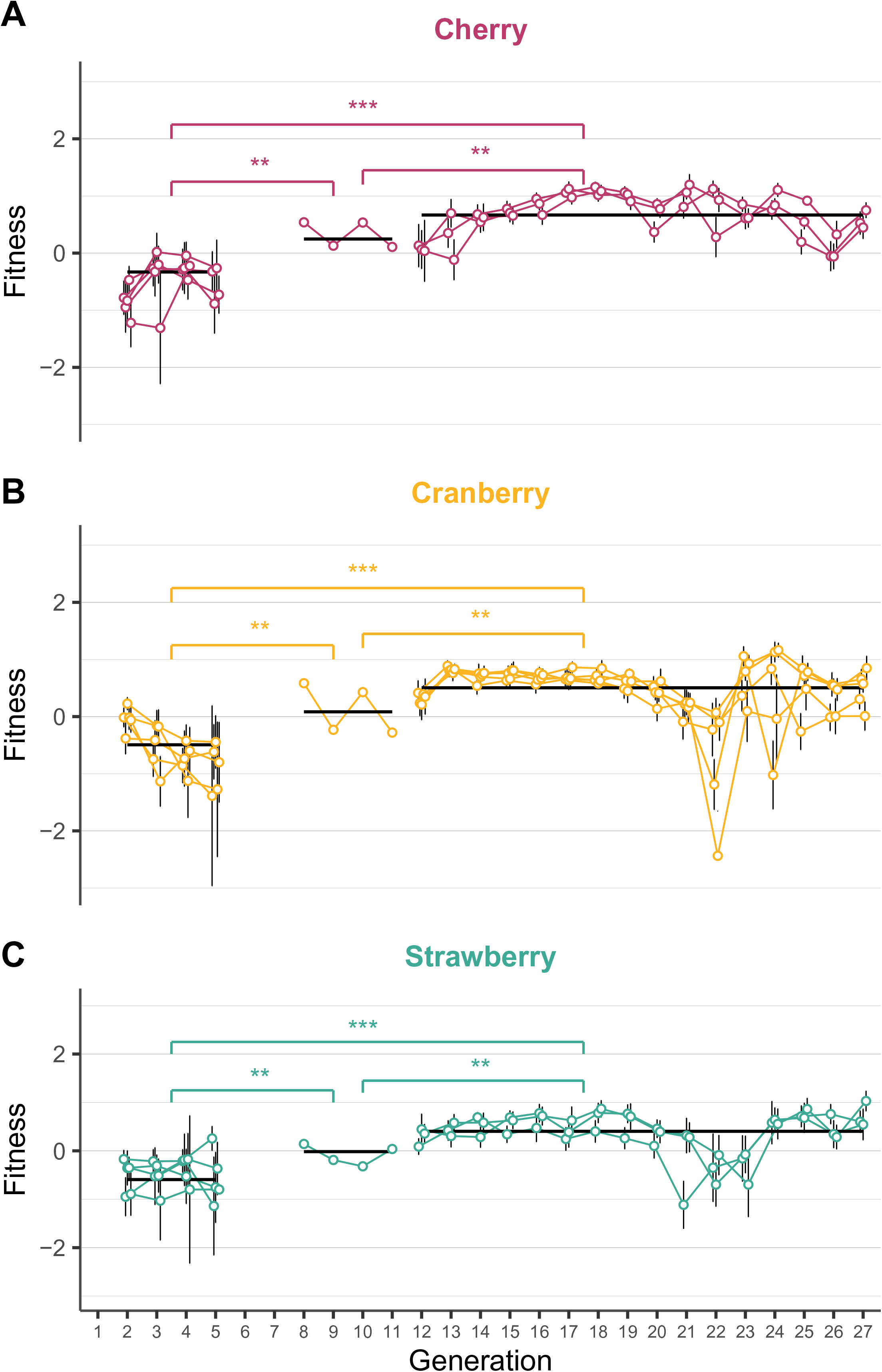
Temporal dynamics of the mean fitness of populations evolved on (A) cherry, (B) cranberry or (C) strawberry during the three phases of the experimental evolution. Malthusian fitness (solid line) was estimated for each of the three phases of the experiment. To avoid the confoundings of maternal effects, data from generations where individuals or their parents developed in standard medium were removed. Error bars represent standard deviation among tubes. The level of significance is provided above horizontal bars (**: *P*-value < 0.01; ***: *P*-value < 0.001).

**Table 2:** Estimate of the change in fitness between the initial and either the intermediate or final phenotyping steps based on the negative binomial model (log scale).

| Phenotyping step | Selective fruit | Test fruit | Estimate | 95% confidence interval |
| --- | --- | --- | --- | --- |
| Intermediate | Cherry | Cherry | -0.02 | [ -0.44 ; 0.37 ] |
|  |  | Cranberry | -0.17 | [ -0.61 ; 0.24 ] |
|  |  | Strawberry | 0.15 | [ -0.32 ; 0.54 ] |
|  | Cranberry | Cherry | -0.01 | [ -0.46 ; 0.42 ] |
|  |  | Cranberry | 0.18 | [ -0.26 ; 0.52 ] |
|  |  | Strawberry | 0.25 | [ -0.17 ; 0.61 ] |
|  | Strawberry | Cherry | 0.46 | [ 0.06 ; 0.81 ] |
|  |  | Cranberry | 0.27 | [ -0.08 ; 0.65 ] |
|  |  | Strawberry | 0.33 | [ -0.02 ; 0.68 ] |
| Final | Cherry | Cherry | 0.26 | [ -0.12 ; 0.6 ] |
|  |  | Cranberry | -0.33 | [ -0.68 ; 0.05 ] |
|  |  | Strawberry | 0.03 | [ -0.37 ; 0.38 ] |
|  | Cranberry | Cherry | 0.25 | [ -0.11 ; 0.6 ] |
|  |  | Cranberry | 0.18 | [ -0.16 ; 0.52 ] |
|  |  | Strawberry | -0.38 | [ -0.75 ; -0.06 ] |
|  | Strawberry | Cherry | 0.02 | [ -0.36 ; 0.41 ] |
|  |  | Cranberry | 0.12 | [ -0.24 ; 0.51 ] |
|  |  | Strawberry | 0.37 | [ -0.07 ; 0.74 ] |
| Overall variance among replicate populations |  |  | 0.14 | [ 0.01 ; 0.17 ] |

Between the start and the end of the experiment (generations 2 and 27 respectively), the average number of adult flies emerging from each tube increased from 8.8 to 35.8, from 19.4 to 33.6 and from 12.2 to 42.3 for cherry, cranberry and strawberry respectively. The corresponding increase observed between phases 1 and 2 were 123%, 93% and 45% and those observed between phases 1 and 3 were 230%, 176% and 145% for cherry, cranberry and strawberry respectively. This increase in fitness across phases had high support (ΔAICc > 27.6 for models 4 to 8 without this effect, Table 1). Support for fitness changes across the three fruit media was strong, while support for differences in temporal fitness increase among fruit media was low (ΔAICc = 3.44 for the model 2 without a fruit effect and ΔAICc = 5.85 for the model 3 with an interaction between the phase and fruit effects, Table 1), indicating that adaptation rates were similar across fruit media. During phase 3, support for fitness changes among populations with each fruit medium was not very strong (ΔAICc = 1.69 for the model 3 including a population effect, Table S4), suggesting that variation in adaptation rate among populations evolving on the same fruit medium tended to be low.

### Changes in fecundity, egg-to-adult viability and fitness in selective and alternative media

Average fecundity increased on the selective environments between the initial and intermediate phenotyping (Table S5, Fig. S3A top panel). In contrast, in egg-to-adult viability were unchanged or decreased modestly between the initial and intermediate phenotyping step, Table S6, Fig. S3B top panel). Changes in fecundity and in egg-to-adult viability were negatively correlated (Fig. S6A). Fitness tended to increase between the first and intermediate genotyping step, with a significant increase for populations that evolved on strawberry and were measured on cherry (large confidence intervals are likely due to a lack of statistical power, Table 2, Fig. 4).

**Figure 4.**
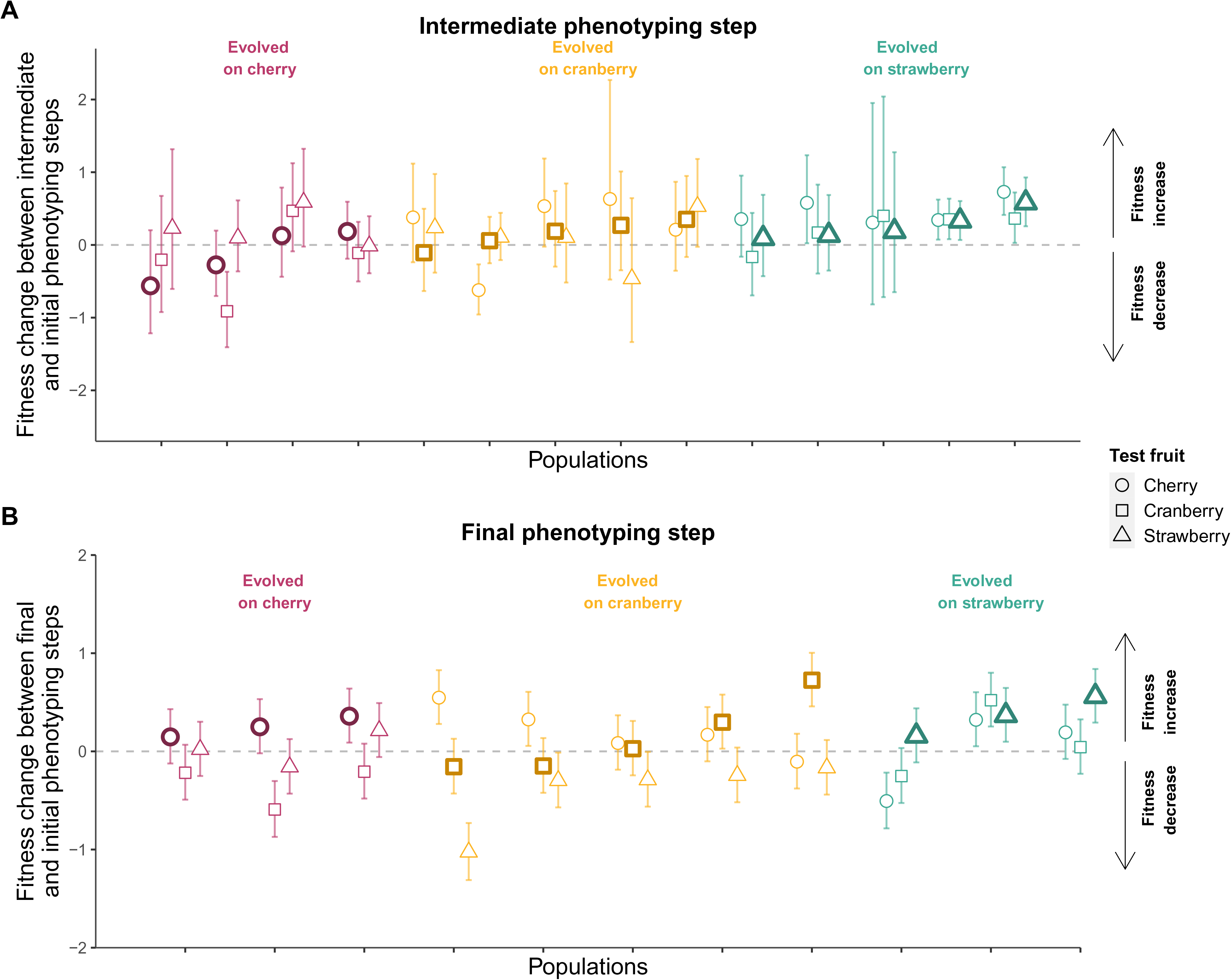
Change in fitness (A) between the initial and intermediate phenotyping steps and (B) between the initial and final phenotyping steps for each population x environment combination. Replicate populations are ordered following their fitness change on selective fruit medium (symbols with thick outline). The color indicates the selective environment, as shown at the top of each group of populations. The shape of the symbol indicates the test fruit as shown in the key to the right. Error bars represent 95% confidence intervals.

Between the initial and final phenotyping steps, no consistent pattern was apparent in average fecundity on the three selective fruits (Table S5, Fig. S3A bottom panel). Egg-to-adult viability increased for populations measured on cherry medium, but did not change consistently or significantly on other media (Table S6, Fig. S4A bottom panel). Changes in fecundity and in egg-to-adult viability were negatively correlated as for the first time-step comparison (Fig. S6B bottom panel). Fitness tended to increase, but not significantly so (Table 2, Fig. 4).

### Correlation between fitness changes in selective and alternative environments

During the intermediate phenotyping step, the increase in fitness in each selective fruit medium was associated with an increase in fitness in the two other fruit media (Fig. 5A-C). This pattern was also seen for some fruit combinations during the final phenotyping step: increases in fitness in selective fruit media were associated with increases in fitness in other fruit media for the pairs strawberry/cranberry and strawberry/cherry (Fig. 5E-F). The confidence intervals of the difference in correlation coefficients between the intermediate and final phenotyping steps overlapped with zero, indicating that the correlation coefficients did not differ significantly. In contrast, increased fitness on cherry or cranberry selective medium were associated with decreased fitness in the reciprocal medium (Fig. 5D). The confidence interval of the difference in correlation coefficients between the intermediate and final phenotyping steps was positive and did not overlap with zero, indicating that the correlation coefficient during the final phenotyping step was lower than that during the intermediate phenotyping step (Fig. 5D). These results remained unchanged when considering only populations that evolved on cranberry (estimate of correlation difference=1.21 [0.32, 1.86]), but not when considering only populations that evolved on cherry medium (estimate of correlation difference=0.55 [-0.71, 1.67]), probably due to a lack of power (see Discussion). The negative correlation in fitness change between cherry and cranberry media was primarily driven by changes in egg-to-adult viability in populations that evolved in cranberry medium, and not by changes in their fecundity (Fig. S4-S5).

**Figure 5.**
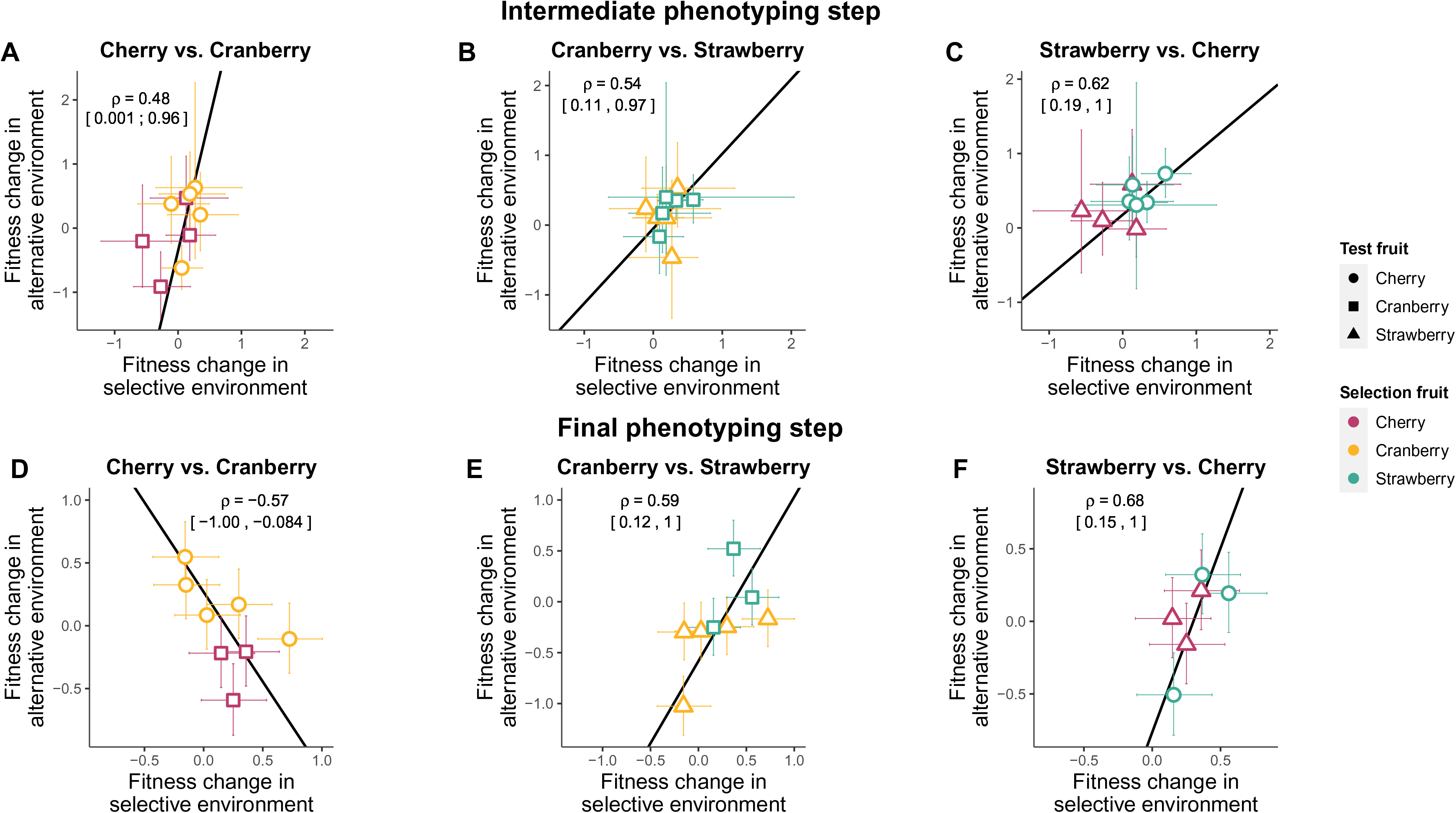
Relationship between the changes in fitness on selective and alternative fruit media between the intermediate and initial phenotyping step (A, B and C) and the final and initial phenotyping step (D, E and F). Solid lines represent fitted major axis slopes. Error bars represent 95% confidence intervals. Correlation coefficients are represented by *ρ* symbols. Values in brackets show 95% confidence intervals. Difference in correlation coefficients between the intermediate and final phenotyping steps for cherry vs. cranberry: 1.05 [0.37, 1.69], cranberry vs. strawberry: -0.06 [-0.65, 0.58] and strawberry vs. cherry: -0.06 [-0.60, 0.60].

Finally, we did not detect a significant pattern of local adaptation during either the intermediate or the final phenotyping steps (*F*_1,3_ < 0.01, *P* = 0.97 and *F*_1,3_ = 2.89, *P* = 0.19 respectively), indicating that fitness changes in selective fruit media were not significantly greater than fitness changes in alternative fruit media.

## Discussion

In the present study, we aimed at quantifying fitness changes in selective and alternative environments of *D. suzukii* experimental populations evolving on each of eight different selective fruit media. Due to the almost complete extinction of populations on five fruit media, we could only estimate fitness changes for populations evolving on cherry, cranberry or strawberry media. After five generations, fecundity had consistently increased in both selective and alternative media, resulting in positive fitness changes across the three fruit media. After 26 generations, both fecundity and egg-to-adult viability had changed compared to the ancestral population. Adaptation to each selective medium was associated with an increase in fitness in alternative media, except for populations that evolved on cranberry when measured on cherry (we had low power to detect the same effect in populations that evolved on cherry medium when measured on cranberry medium). Indeed, egg-to-adult viability on cherry and cranberry media were negatively correlated for populations that evolved on cranberry. These results suggest that cranberry and cherry media might exert very different selective pressures on egg-to-adult viability.

### Relative importance of selection, genetic drift and pleiotropy in shaping fitness changes in selective and alternative environments

During the first five generations of evolution in phase 1, we found weak evidence for the adaptation of populations evolving on each of the eight fruit media, which could suggest the absence of genetic variation in adaptive alleles or the absence of selection (i.e., that adaptive alleles did not increase in frequency in these populations). This interpretation might be overly simplistic for two reasons. First, Olazcuaga (2019) recently found that local adaptation to different fruit occurs over less than four generations in natural populations, which suggests that genetic variation in adaptive alleles present in natural populations likely persisted in our lab population. Second, we observed that the fitness of experimental populations evolving on cherry, cranberry and strawberry media increased significantly following the pooling step and in the subsequent generations (Fig. 3), which indicates that natural selection is present in our experiment.

Furthermore, two different processes could account for the increase in fitness after the pooling step. First, the increased fitness might be due to heterosis following the masking of mildly deleterious alleles that independently increase in frequency in replicate populations during phase 1 (low population sizes likely increased genetic drift during phase 1 so that mildly deleterious mutations became selectively neutral). Second, the increase in fitness might be due to the combination of different adaptive alleles that independently increase in frequency in replicate populations evolving on the same fruit medium during phase 1 (Barghi et al. 2019). Note, that the occurrence of only one of the two processes during phase 1 would likely have resulted in a fitness change (a decrease in fitness with increase in frequency of deleterious alleles or an increase in fitness with an increase in adaptive alleles). As we did not detect such a change, both processes likely occurred concurrently during phase 1; the negative fitness effect associated with the increase in frequency of mildly deleterious alleles across the genome might have been counterbalanced by the positive fitness effects associated with the increase in frequency of a few large-effect mutations, as observed in Stewart et al. (2017) and Koch and Guillaume (2020). Finally, we did not detect a pattern of local adaptation during the intermediate phenotyping step, which suggests that beneficial mutations that increased in frequency might not have been fruit-specific. However, this result might also be due to our low statistical power, so that this interpretation remains speculative. Genomic data could help in estimating the extent of genetic drift.

During phase 3, the fitness changes of populations evolving on the same fruit medium tended to be in the same direction, while their fitness changes in each of the two other fruit media also tended to be in the same direction. Fitness changing in a consistent way across replicate populations is likely the result of natural selection, as genetic drift would result in the evolution of fitness increasing or decreasing at random across replicate populations. Furthermore, the convergent evolution of fitness changes in alternative fruit media suggests that the same trait(s) might have been selected independently in replicate populations adapting to the same fruit medium. An alternative explanation would be that this pattern arose through the independent evolution of different phenotypic traits in replicate populations (due to selection or to genetic drift). This would require these different traits to have the exact same pleiotropic fitness effects across fruit media. The precise relationship between a given phenotypic trait and fitness is often environment-specific, so that this alternative hypothesis appears quite unlikely. Genomic data would be necessary to thoroughly assess and quantify convergence among replicate populations at the molecular level (e.g., following Barghi et al. 2019).

Finally, the demographic trajectories we observed after the temporary pooling suggest the occurrence of evolutionary or genetic rescue (Gomulkiewicz and Holt 1995; Whiteley et al. 2015; Hedrick and Garcia-Dorado 2016). However, although the data do fit that interpretation well, our experiment was not set up explicitly to study this phenomenon, and we lack the appropriate experimental controls to confirm or infirm this interpretation.

### Fitness in selective and alternative environments shed light on the hypothetical fitness landscape

The main focus of our study was to use distinct environments to test for a reversal over time in the direction of the association between fitness changes in selective and alternative environments. Such a reversal was observed for populations that evolved on cranberry when measured on cherry, confirming our predictions based on fitness landscapes theory and Fisher’s geometric model (Martin & Lenormand 2015).

More generally, our results can be used in an interpretation of the theoretical fitness landscape represented by different environments. The increase in fitness in both selective and alternative environments observed during the intermediate phenotyping step clearly indicates that the ancestral population was initially maladapted to each of the three selective fruit media in a similar way (Figs. 1 & 6). Alleles favored early in the experiment likely increased fitness across the three selective media, as adaptation occurred mostly through an increase in fecundity that was of similar magnitude across replicate populations and across fruit media (Fig. S3A). Given that the initial rate of adaptation was similar across the three fruit media, the distance between the ancestral population and the phenotypic optimum of each of the three fruit media was likely similar (this interpretation parsimoniously assumes that the levels of adaptive genetic variation to each of the three fruit media were similar in the ancestral population and that the intensity of selection was also similar across fruit media). Adaptation occurred through changes in both fecundity and egg-to-adult viability that depended on the population or on the selective fruit (Fig. S3). Eventually, populations that evolved on cranberry, exhibited reduced fitness in one of the alternate environments (cherry), which suggests that the phenotypic optima of these two fruit media lie the furthest away from each other in the fitness landscape (Fig. 6).

**Figure 6.**
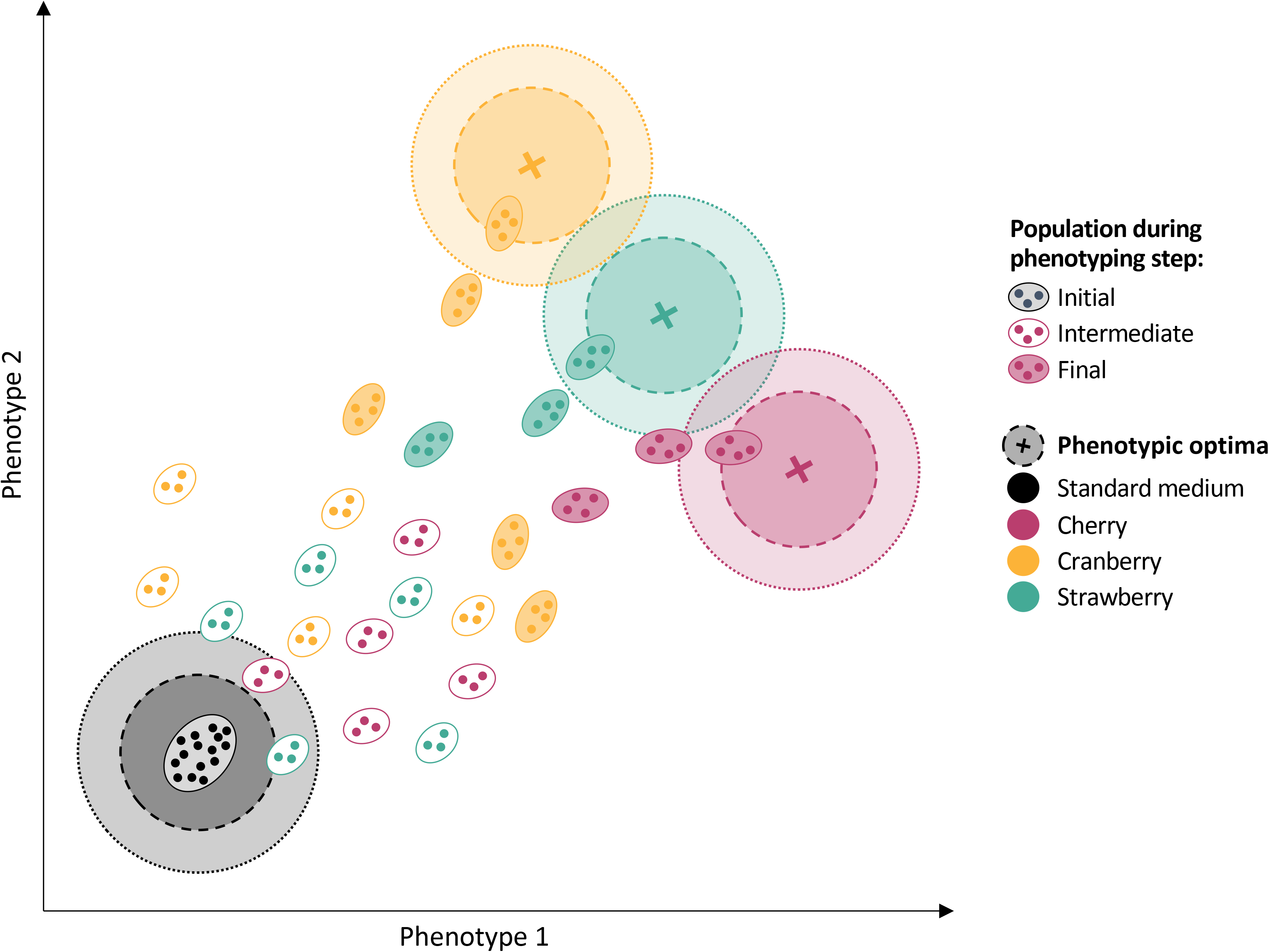
Hypothetical fitness landscape to help in the interpretation of our results. For each environment, the position of the phenotypic optimum providing maximal fitness is represented by a cross. The positions of populations and genotypes within populations are respectively represented by ellipses and closed circles. We hypothesized that adaptation during phase 1 was masked by the increase in frequency of mildly deleterious mutations due to genetic drift (see Discussion), populations during the intermediate phenotyping step are represented closer to the fruit media optima than the ancestral population, although no significant fitness change was detected.

This pattern was primarily driven by changes in egg-to-adult viability in populations that evolved in cranberry. We had low power to detect similar changes in populations that evolved in the cherry medium. This example (Fig. 6) illustrates how examining direct and correlated responses to selection over time can help visualize the fitness landscapes organisms experience in different environments.

### Limits of our experimental approach

The fitness optima of the five fruit media where populations went extinct were likely further away than the fitness optima of the three fruit media where populations survived. Consequently, we could not assess fitness changes across fruit environments that exert the most divergent selective pressures (i.e., cherry, cranberry, and strawberry vs. the five other fruits). Hence, population extinctions have limited our power to detect a reversal in fitness correlations among fruit media. Despite the likely closer proximity of the fitness optima of the three remaining fruit media (cherry, cranberry, and strawberry), we nevertheless detected a reversal in the fitness correlation between cherry and cranberry media.

Fitness changes could be caused by temporal variation in environmental conditions in the laboratory (e.g., change of quality of frozen fruit purees). However, temporal environmental variation would be unlikely to result in adaptation to selective media and thus unlikely to explain our findings.

While the temporary pooling of replicate populations most likely facilitated further adaptation by reducing inbreeding depression, it might have limited our power of inference regarding the variation in fitness changes among replicate populations. Indeed, changes in fitness of replicate populations in each selective medium or in the two alternative media tended to be in the same direction during phase 3. This pattern might be due to the co-ancestry of replicate populations. In other words, the observed fitness changes might have evolved in a single replicate population during phase 1 or phase 2, rather than several times independently in each replicate population during the 16 later generations. We consider this hypothesis to be unlikely for four reasons. First, the fitness of replicate populations evolving on each fruit increased significantly between the intermediate and final phenotyping steps, when populations evolved independently. This demonstrates that the observed fitness changes are partly independent of each other. Second, at the end of phase 3, fitness changes in selective or alternative environments varied among replicate populations evolving on the same fruit medium, which supports the view that fitness changes are partially independent. Third, the simulations of unequal contributions of replicate populations to the pool during phase 2 show that the pattern of reversal in the association between fitness changes in selective and alternative environments (Fig. 5D) cannot be explained by the pooling step (see Appendix S4, Fig. S10 and S11). Fourth, the effect of pooling on reducing inbreeding depression is probably stronger than its effect on increasing the frequency of adaptive mutations. Indeed, we expect the five generations of pooling would mask the effect of the numerous deleterious alleles located all over the genome but would not favor the spread of beneficial alleles confined to particular loci within the genome.

### Recommendations for using fitness landscapes to interpret selection experiments

We can make several recommendations for future studies that aim to track and predict the evolutionary trajectories of experimental populations evolving in contrasted environments. First, measuring fitness rather than fitness proxies allows for a standard comparison among populations reared on different (i.e., selective and alternative) environments. For example, fitness can be used to compare populations evolving with different maintenance schemes (for example, egg-to-adult viability over two weeks versus two months). Second, when using experimental evolution to study fitness changes in selective and alternative environments, the likely position of the ancestral population in the fitness landscape relative to the phenotypic optima of selective environments should be considered with care. For example, many studies of ecological specialization using experimental evolution find positive instead of negative fitness correlated responses in alternative environments (Futuyma 2008). As explained in the introduction, these results can be explained by two alternative and mutually exclusive hypotheses (Fig. 1). On the one hand, if the environments have the same fitness optima, additional generations of experimental evolution would still result in positive correlation in fitness across environments (Fig. 1A). On the other hand, if the environments have different fitness optima, the level of maladaptation of the ancestral population matters. When starting from an ancestral population similarly maladapted to the two environments, additional generations of experimental evolution would result in negative fitness correlation across environments (Fig. 1B), as illustrated in our study. Following other studies (e.g., Fragata et al. 2019), we emphasize that fitness landscape theory represents a powerful framework for studying the process of local adaptation using experimental evolution. Third, how long it takes for experimental evolution to show a reversal in the direction of fitness changes across environments depends on the level of adaptive genetic variation in the ancestral population. Negative correlations in fitness can evolve over short time scales when initial levels of standing adaptive genetic variation are high, as exemplified in our study. In contrast, negative correlations in fitness likely evolve more slowly in studies based on de novo mutation, as illustrated by Bono et al 2017.

### Conclusion and perspectives

We found temporal adaptation in *D. suzukii* experimental populations evolving in three different selective environments. Adaptation to each fruit was associated with an increase in fitness in the two other fruits with the exception of populations that evolved either on cherry or on cranberry medium. Our results show that the temporal study of fitness changes in selective and alternative environments across multiple generations allows a better characterization of the dynamics of local adaptation compared to typical cross-sectional studies performed over a single generation. This framework could improve our understanding of the ecological factors that drive the evolution of local adaptation.

## Data availability statement

Data code for our analyses are available at Dryad https://doi.org/10.5061/dryad.crjdfn33t

## Acknowledgements

We are grateful to A. Savage, A. Gagu, L. Benoit, J.L. Imbert and R. Vedovato for technical assistance, S. Magalhães, T. Guillemaud, T. Lenormand, L.-M. Chevin, G. Martin and E. Lievens for comments on the manuscript and for insightful discussions, and the Sicoly cooperative for providing us with several of the fruit purees. We are grateful to the SEPA technical platform of the CBGP laboratory for hosting all experiments presented in this study. L.O., M.G., J.F. and A.E. were supported by the Languedoc-Roussillon region (France) through the European Union program FEFER FSE IEJ 2014-2020 (project CPADROL) and the INRAE scientific department SPE (AAP-SPE 2016), and the French Agence National pour la Recherche (ANR-16-CE02-0015). R.A.H. acknowledges support from the National Science Foundation (DEB-0949619), USDA Agriculture and Food Research Initiative award (2014- 67013-21594), Hatch project 1012868, the French Agropolis Fondation (LabEx Agro– Montpellier) through the AAP “International Mobility” (CfP 2015-02), the French programme investissement d’avenir, and the LabEx CEMEB through the AAP “invited scientist 2016”. L.O. and N.O.R. acknowledge support from the CeMEB LabEx/University of Montpellier (ANR-10-LABX-04-01). V.R. received support from the European Regional Development Fund (ERDF) and the Conseil Régional de la Réunion. We are grateful to the SEPA technical platform of the CBGP laboratory for hosting all experiments presented in this study.

## Author contributions

Conceptualization, L.O., N.O.R., J.F., M.G., B.F., V.R., R.A.H. and A.E.; Experimental design, L.O., N.O.R., J.F. and R.A.H.; Experiment realization and data acquisition, L.O., C.D., A.L. and N.L.; Statistical analysis, L.O. and N.O.R.; Writing – Original Draft, L.O. and N.O.R.; Writing – Review & Editing, J.F., M.G., B.F., V.R., R.A.H. and A.E.; Funding Acquisition, M.G. and A.E.

## Conflict of interest statement

The authors have no conflict of interest to declare.

## Notes

### Competing Interest Statement

The authors have declared no competing interest.

