## Supplementary materials for "Adaptation and correlated fitness responses over two time scales in *Drosophila suzukii* populations evolving in different environments"

#### Table of content

**Supplementary Table**
Table S1

**Appendix S1**
Tables S2-S4
Figure S1

**Appendix S2**
Tables S5-S6
Figure S2-S6

**Appendix S3**
Figure S7-S9

**Appendix S4**
Figure S10-S11

### Supplementary Table

##### Table S1. Potential host-plant used by *D. suzukii* during the sampling period

List of host plants used by *D. suzukii* (based on fruits reported as host in the field; Kenis et al. 2016) that could have been present in the sampled field area during the sampling period (presence and period of fruiting period was obtained using the open access national naturalist networks tela-botanica.org).

| Host-plant species | Family | Fruiting period |
|---|---|---|
| *Actinidia chinensis* | Actinidiaceae | September |
| *Amelanchier ovalis* | Rosaceae | August-September |
| *Arbutus unedo* | Ericaceae | October-January |
| *Cornus mas* | Cornaceae | September |
| *Cornus sanguinea* | Cornaceae | September-October |
| *Lycium barbarum* | Solanaceae | March-October |
| *Phytolacca americana* | Phytolaccaceae | June-September |
| *Pyracantha sp.* | Rosaceae | September-October |
| *Rhamnus cathartica* | Rhamnaceae | August-September |
| *Sambucus ebulus* | Adoxaceae | September-October |
| *Sambucus nigra* | Adoxaceae | September |
| *Sambucus racemosa* | Adoxaceae | July-September |
| *Solanum dulcamara* | Solanaceae | June-September |
| *Solanum nigrum* | Solanaceae | June-November |
| *Sorbus aria* | Rosaceae | September |
| *Sorbus aucuparia* | Rosaceae | September |
| *Taxus baccata* | Taxaceae | August-September |
| *Viburnum lantana* | Adoxaceae | August-September |

### Appendix S1: Heterogeneity in fitness changes among populations during phases 1 and 3

#### Methods

To test for differences in the rate of adaptation among fruit media and among populations during either phase 1 or phase 3, we fitted the following linear model on fitness $m_{ijk}$:

$$m_{ijk} = generation_i + population_j + selective\_fruit_k + generation{:}population_{ij} + generation{:}selective\_fruit_{ik} + \underline{generation{:}selective\_fruit_{ij}} + \varepsilon_{ijk} \quad (4),$$

where fixed effects included the effect of the *i*th generation (*generation$_i$*, with *i*=2, ..., 5 for phase 1 or *i*=12, ..., 27 for phase 3), the effect of the *j*th population (*population$_j$*, with *k*=1, ..., 40 for phase 1 or *k*=1, ..., 11 for phase 3), the effect of the *k*th selective fruit (*selective_fruit$_k$*, with *k*=1,..,8 for cherry, strawberry, cranberry, blackcurrant, fig, grape, rose hips, and tomato respectively), the interaction between the *generation* and *population* effects, the interaction between the *generation* and *selective_fruit* effects - where random effects included the interaction between the effect of the *i*th generation and the effect of the *j*th *selective_fruit* (mean of zero and variance $\sigma^2_{gen\ fruit}$), and a random error ($\varepsilon_{ij}$ mean of zero and variance $\sigma^2_{res}$). Due to their redundancy, the *population* and *selective_fruit* effects were never tested together. We used the same AICc model selection as in the main text.

#### Results

During phase 1, fitness estimates for each of the eight fruit media were negative (Fig. S1). We found high support for fitness changes across fruit media (ΔAICc > 28.3 for models 14, 15 and 16 without a fruit effect, Table S2), but not across populations (ΔAICc = 9.48 for the model 12 including a population effect, Table S2). Temporal changes in fitness differed across fruits

(ΔAICc = 2.12 for the model 10 with neither a generation effect nor an interaction between fruit and generation effects, Table S2).

During phase 3, we found high support for variation in fitness changes across selective fruit media (ΔAICc > 2.5 for models 20 to 24 without a fruit effect, Table S4), but not for variation across populations within each fruit medium (ΔAICc = 1.69 for the model 19 including a population effect, Table S4). Finally, we found no support for a temporal increase in fitness in the different fruit media (the model 18 including both a generation and a fruit effect had almost the same log-likelihood as the model 17 with only a fruit effect, Table S4).

**Table S2: Results of the AICc model selection for fitness changes in selective environments during phase 1.** All models included the interaction between generation and selective_fruit as a random effect.

| Models | Effects | df | logLik | AICc | ΔAICc |
|---|---|---|---|---|---|
| (9) | Generation × Selective_fruit | 18 | -80.61 | 205.00 | 0.00 |
| (10) | Selective_fruit | 10 | -92.41 | 207.10 | 2.12 |
| (11) | Generation + Selective_fruit* | 11 | -92.31 | 209.40 | 4.40 |
| (12) | Population | 42 | -37.02 | 214.50 | 9.48 |
| (13) | Generation + Population | 43 | -35.79 | 217.60 | 12.65 |
| (14) | Generation | 4.00 | -112.48 | 233.40 | 28.36 |
| (15) | Null Model | 3 | -114.23 | 234.70 | 29.70 |
| (16) | Generation × Population | 71 | 6.83 | 420.50 | 215.47 |

* This model had the same log-likelihood as a model without a generation effect.

**Table S3: Intercept and generation slope estimates and standard errors (SE) from the model with lowest AICc during phase 1.**

| Fruit media | Intercept | SE intercept | Slope | SE slope |
|---|---|---|---|---|
| Blackcurrant | -1.26 | 1.44 | -0.56 | 0.70 |
| Cherry | -1.12 | 1.58 | 0.18 | 0.76 |
| Cranberry | 0.46 | 1.58 | -0.29 | 0.76 |
| Fig | -0.11 | 1.62 | -0.31 | 0.77 |
| Grape | -4.71 | 3.40 | 1.11 | 1.66 |
| Rosehips | -3.87 | 1.83 | 0.78 | 0.87 |
| Strawberry | -0.70 | 1.58 | 0.08 | 0.76 |
| Tomato | -2.83 | 1.89 | 0.55 | 0.88 |

**Table S4: Results of the AICc model selection for fitness changes in selective environments during phase 3.** All models included the interaction between generation and selective_fruit as a random effect.

| Models | Effects | df | logLik | AICc | ΔAICc |
|---|---|---|---|---|---|
| (17) | Selective_fruit | 5 | -85.57 | 181.50 | 0.00 |
| (18) | Generation + Selective_fruit* | 6 | -85.08 | 182.60 | 1.15 |
| (19) | Population | 13 | -77.47 | 183.20 | 1.69 |
| (20) | Modele Null | 3 | -88.95 | 184.00 | 2.54 |
| (21) | Generation + Population | 14 | -76.90 | 184.40 | 2.92 |
| (22) | Generation | 4 | -88.57 | 185.40 | 3.88 |
| (23) | Generation × Selective_fruit | 8 | -84.56 | 186.00 | 4.48 |
| (24) | Generation × Population | 24 | -67.42 | 190.80 | 9.30 |

* This model had the same log-likelihood as a model without a generation effect.

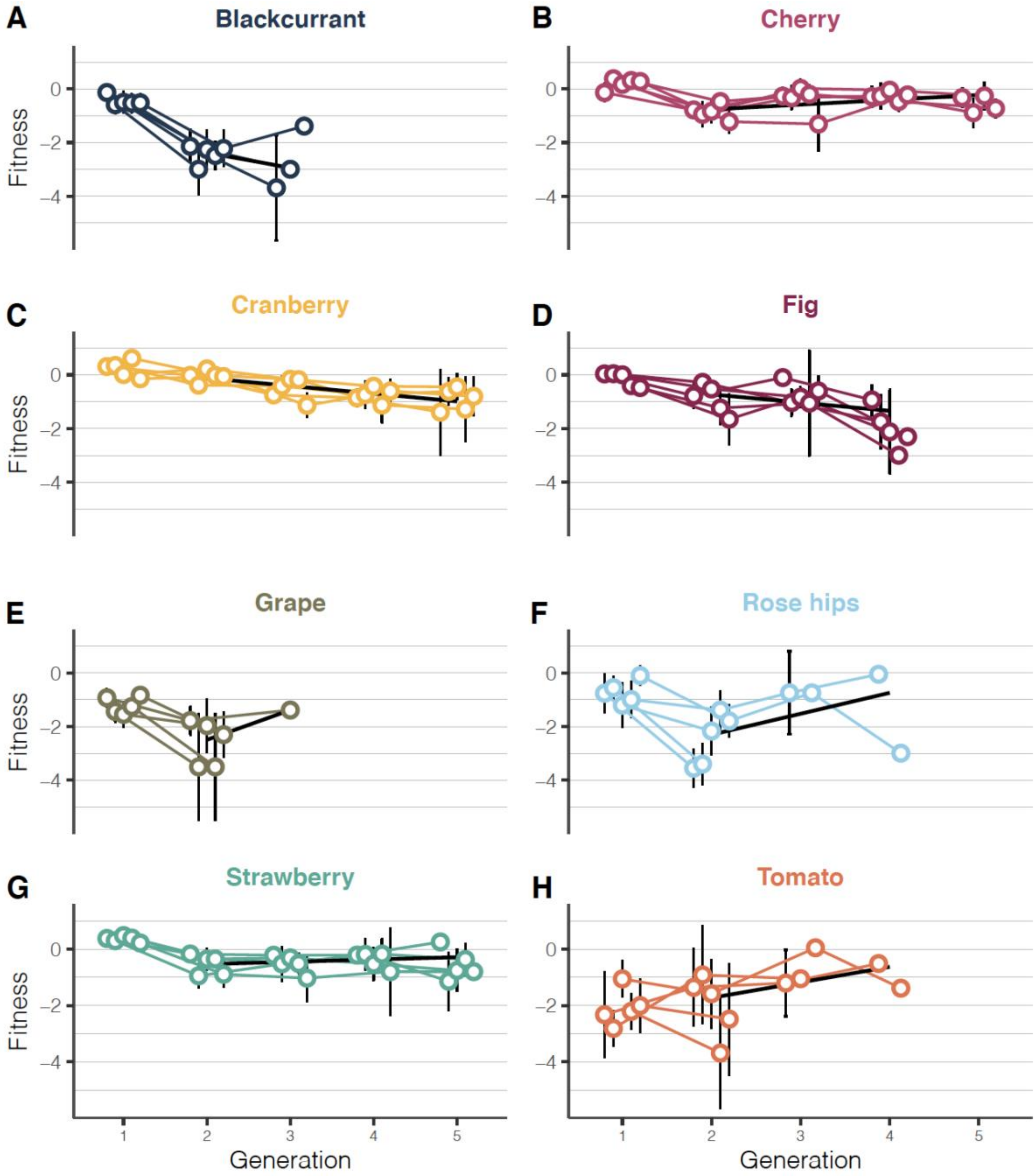


**Figure S1. Temporal dynamics of the mean fitness of populations evolved on (A) blackcurrant, (B) cherry, (C) cranberry, (D) fig, (E) grape, (F) rose hips, (G) strawberry or (H) tomato during the first phase of experimental evolution.** Solid lines represent the fitted Malthusian fitness, excluding the first generation to avoid the confounding maternal effect of having reared the ancestral population on standard medium. The absence of data for later generations in some fruit media was due to population extinction. Error bars represent standard deviation among tubes.

### Appendix S2: Changes in female fecundity, egg-to-adult viability and fitness in selective and alternative media

For each population and for each of the three fruit media, we computed the difference in Malthusian fitness between the phenotyping step *n* and the initial phenotyping step as: $s_n=m_n-m_{initial}$ (Fisher 1930, Chevin 2011). We combined the data of the three phenotyping steps and fitted the following negative binomial model (log link) on $n_{ij}$, that represent either the number of eggs in each tube or the number of adults that emerged from each tube:

$$\log(n_{ij}) = test\_fruit_i + test\_fruit{:}population_{ij} \quad (5),$$

where $test\_fruit_i$ represents the effect of the ancestral population on the *i*th test fruit medium ($test\_fruit_i$, with i=1, 2 and 3, for cherry, cranberry and strawberry respectively) and the interaction $test\_fruit{:}population_{ij}$ represents the effect of the *j*th populations step on the *i*th test fruit medium ($population_j$, with either j=1,... 14 and j=15,... 25 for the intermediate and final phenotyping steps, respectively). We used the *glm.nb* function of the *MASS* package and computed the profile likelihood 95% confidence interval of each parameter using the confint function (Venables and Ripley 2013).

For egg-to-adult viability, we fitted the following binomial model (logit link) on $p_{ij}$ (number of adults that emerged out of the total number of eggs counted in each tube:

$$\text{logit}(p_{il}) = test\_fruit_i + test\_fruit{:}population_{ij} \quad (6),$$

were parameters are as those described above. We used the *glm* function of the *stats* package and computed the profile likelihood 95% confidence interval of each parameter using the confint function (Venables and Ripley 2013).

We performed simulations based on our fitness data to verify that the 95% confidence interval of the fitness change between evolved and ancestral populations was accurately estimated by likelihood profiling. For each simulation, we sampled the number of adults emerging from each tube using a negative binomial distribution (*rnegbin* function in the *MASS* package). We used different average numbers of individuals for the ancestral and evolved populations (i.e., a mean of 25 and 30 individuals respectively), but the same overdispersion parameter (theta=3). To mimic our experimental data, we considered that the fitness of the ancestral and evolved populations were estimated based on a different number of vials (n=100 vials for the ancestral population and n varying from 4 to 32 vials for the evolved populations). For each simulation, we computed the average fitness difference between ancestral and evolved populations and its likelihood profile 95% confidence interval. We performed 1000 simulations and estimated the proportion of times where the true fitness difference (i.e log(30/25)) fell within the estimated 95% confidence interval. Our simulations showed that the coverage of the 95% confidence interval was accurate (Fig. S2).

**Table S5: Estimate of fecundity changes between the initial and either the intermediate or final phenotyping steps based on the negative binomial model (log scale).**

| Phenotyping step | Selective fruit | Test fruit | Estimate | 95% confidence interval |
|---|---|---|---|---|
| Intermediate | Cherry | Cherry | 0.27 | [ 0.11 ; 0.41 ] |
| | | Cranberry | 0.35 | [ 0.21 ; 0.5 ] |
| | | Strawberry | 0.48 | [ 0.33 ; 0.65 ] |
| | Cranberry | Cherry | 0.22 | [ 0.08 ; 0.37 ] |
| | | Cranberry | 0.39 | [ 0.28 ; 0.53 ] |
| | | Strawberry | 0.31 | [ 0.17 ; 0.45 ] |
| | Strawberry | Cherry | 0.20 | [ 0.08 ; 0.32 ] |
| | | Cranberry | 0.46 | [ 0.35 ; 0.59 ] |
| | | Strawberry | 0.48 | [ 0.37 ; 0.6 ] |
| Final | Cherry | Cherry | -0.37 | [ -0.47 ; -0.26 ] |
| | | Cranberry | 0.02 | [ -0.09 ; 0.12 ] |
| | | Strawberry | 0.20 | [ 0.09 ; 0.3 ] |
| | Cranberry | Cherry | -0.30 | [ -0.41 ; -0.21 ] |
| | | Cranberry | 0.01 | [ -0.08 ; 0.1 ] |
| | | Strawberry | 0.14 | [ 0.04 ; 0.23 ] |
| | Strawberry | Cherry | -0.47 | [ -0.59 ; -0.37 ] |
| | | Cranberry | -0.13 | [ -0.23 ; -0.01 ] |
| | | Strawberry | 0.16 | [ 0.05 ; 0.27 ] |
| Overall variance among replicate populations | | | 0.00 | [ 0 ; 0.02 ] |

**Table S6: Estimate of changes in egg-to-adult viability between the initial and either the intermediate or final phenotyping steps based on the binomial model (logit scale).**

| Phenotyping step | Selective fruit | Test fruit | Estimate | 95% confidence interval |
|---|---|---|---|---|
| Intermediate | Cherry | Cherry | -0.31 | [ -0.62 ; -0.02 ] |
| | | Cranberry | -0.53 | [ -0.84 ; -0.25 ] |
| | | Strawberry | -0.38 | [ -0.67 ; -0.09 ] |
| | Cranberry | Cherry | -0.10 | [ -0.4 ; 0.16 ] |
| | | Cranberry | -0.14 | [ -0.42 ; 0.12 ] |
| | | Strawberry | 0.02 | [ -0.27 ; 0.28 ] |
| | Strawberry | Cherry | 0.24 | [ -0.02 ; 0.53 ] |
| | | Cranberry | -0.22 | [ -0.5 ; 0.08 ] |
| | | Strawberry | -0.17 | [ -0.45 ; 0.11 ] |
| Final | Cherry | Cherry | 0.62 | [ 0.34 ; 0.93 ] |
| | | Cranberry | -0.35 | [ -0.64 ; -0.05 ] |
| | | Strawberry | -0.17 | [ -0.45 ; 0.14 ] |
| | Cranberry | Cherry | 0.54 | [ 0.29 ; 0.78 ] |
| | | Cranberry | 0.19 | [ -0.06 ; 0.45 ] |
| | | Strawberry | -0.52 | [ -0.77 ; -0.27 ] |
| | Strawberry | Cherry | 0.52 | [ 0.25 ; 0.84 ] |
| | | Cranberry | 0.25 | [ -0.01 ; 0.54 ] |
| | | Strawberry | 0.18 | [ -0.1 ; 0.46 ] |
| Overall variance among replicate populations | | | 0.18 | [ 0.11 ; 0.22 ] |

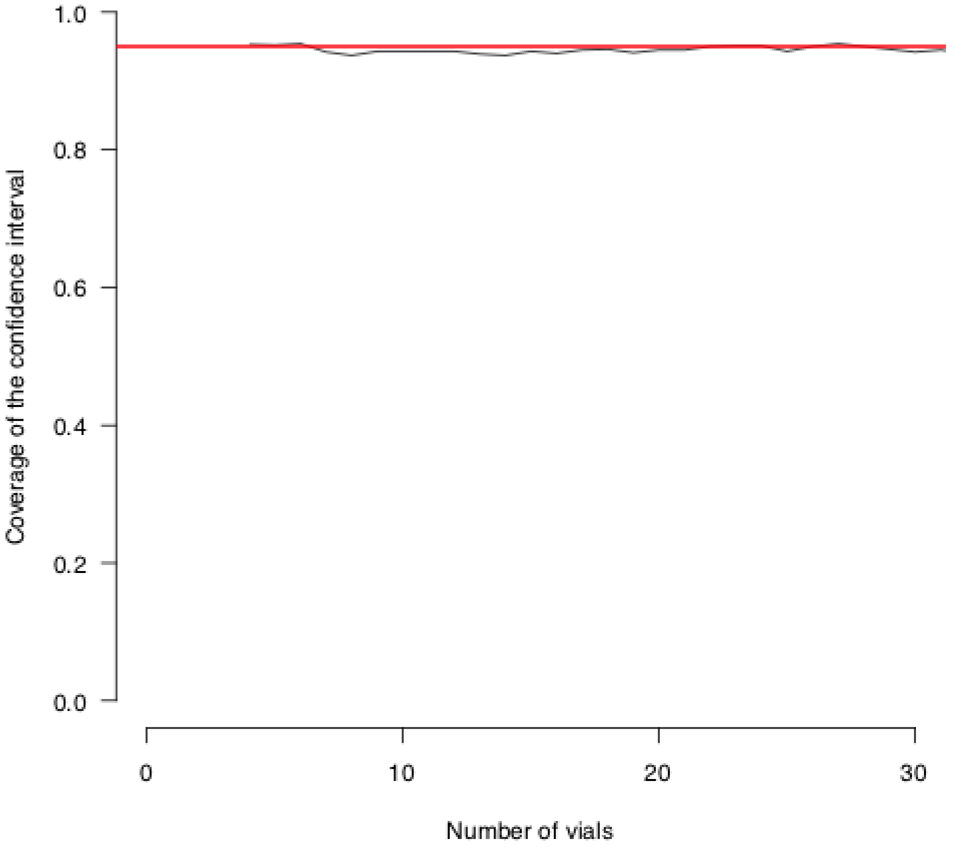


**Figure S2. Coverage of the confidence interval of the fitness difference between ancestral and evolved populations as a function of the number of vials used for the evolved population.The thin black line represents coverage estimates based on 10000 simulated datasets. The red horizontal line represents a 95% threshold coverage.**

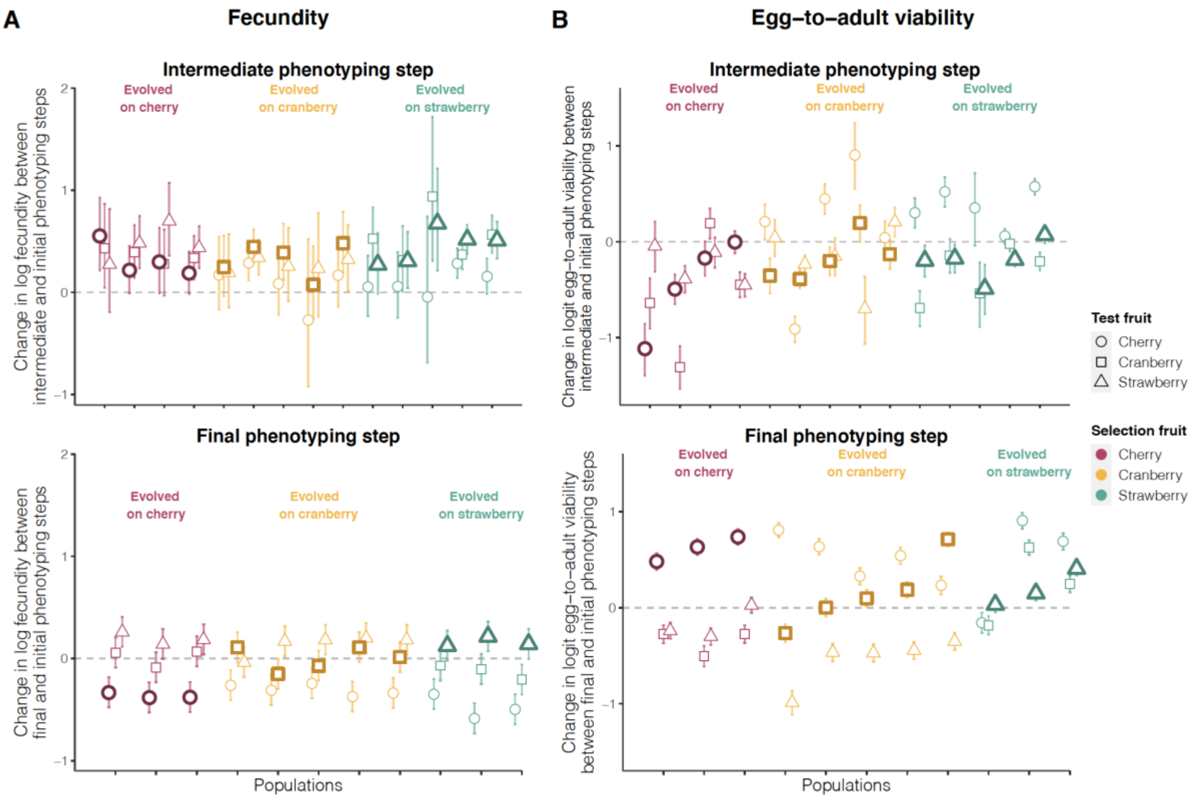


**Figure S3. Change in (A) fecundity and (B) egg-to-adult viability between the initial and intermediate phenotyping steps (top panel) and between the initial and final phenotyping steps (bottom panel) for each population x environment combination**. Replicate populations are ordered following their fitness change on selective fruit medium (symbols with thick outline). The color indicates the selective environment, as shown at the top of each group of populations. The shape of the symbol indicates the test fruit as shown in the key to the right. Error bars represent 95% confidence intervals.

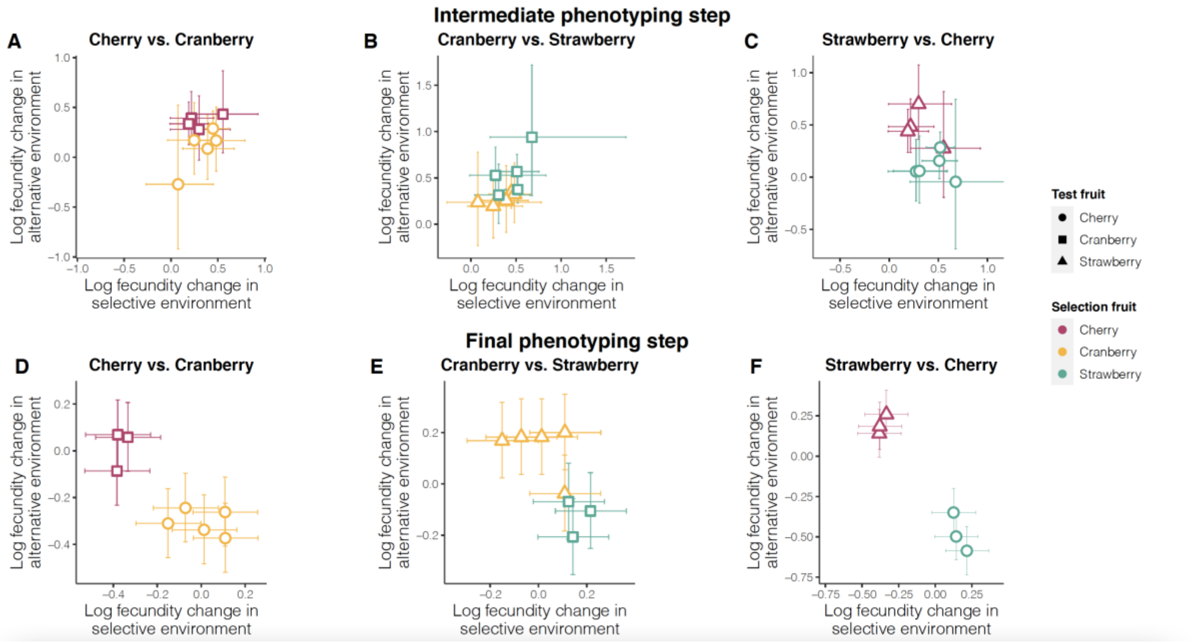


**Figure S4. Relationship between the fecundity changes on selective and alternative fruit media between either the intermediate (A, B and C) or the final phenotyping steps (D, E and F) and the initial phenotyping step.** Error bars represent 95% confidence intervals.

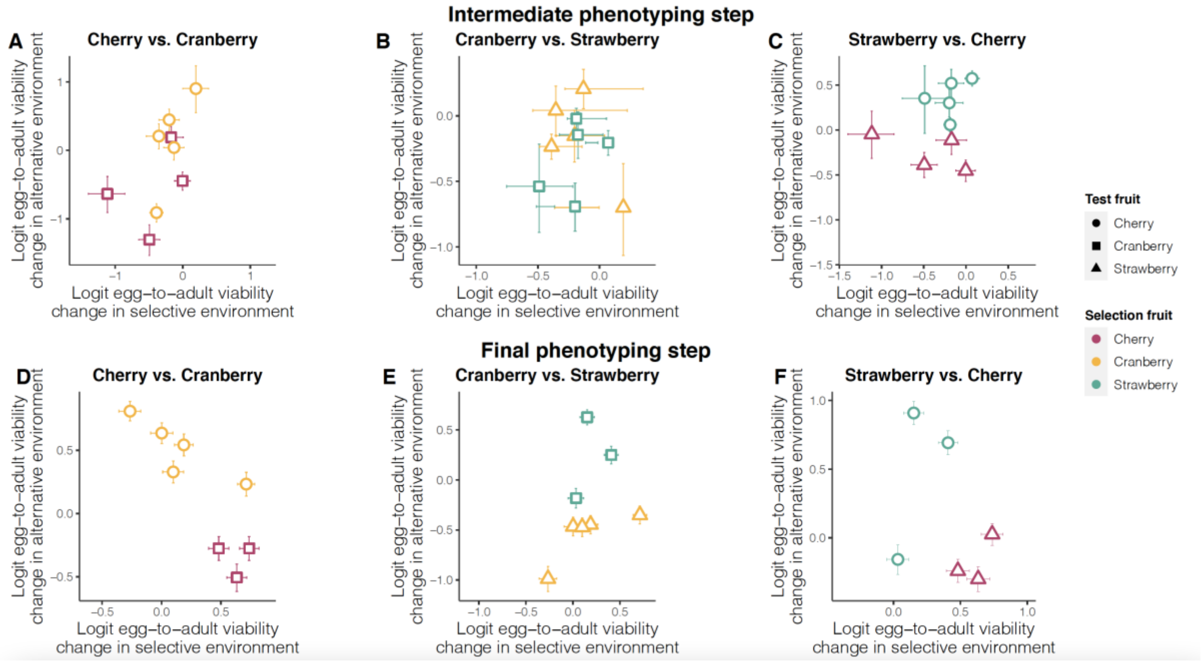


**Figure S5. Relationship between the changes in egg-to-adult viability on selective and alternative fruit media between either the intermediate (A, B and C) or the final phenotyping steps (D, E and F) and the initial phenotyping step.** Error bars represent 95% confidence intervals.

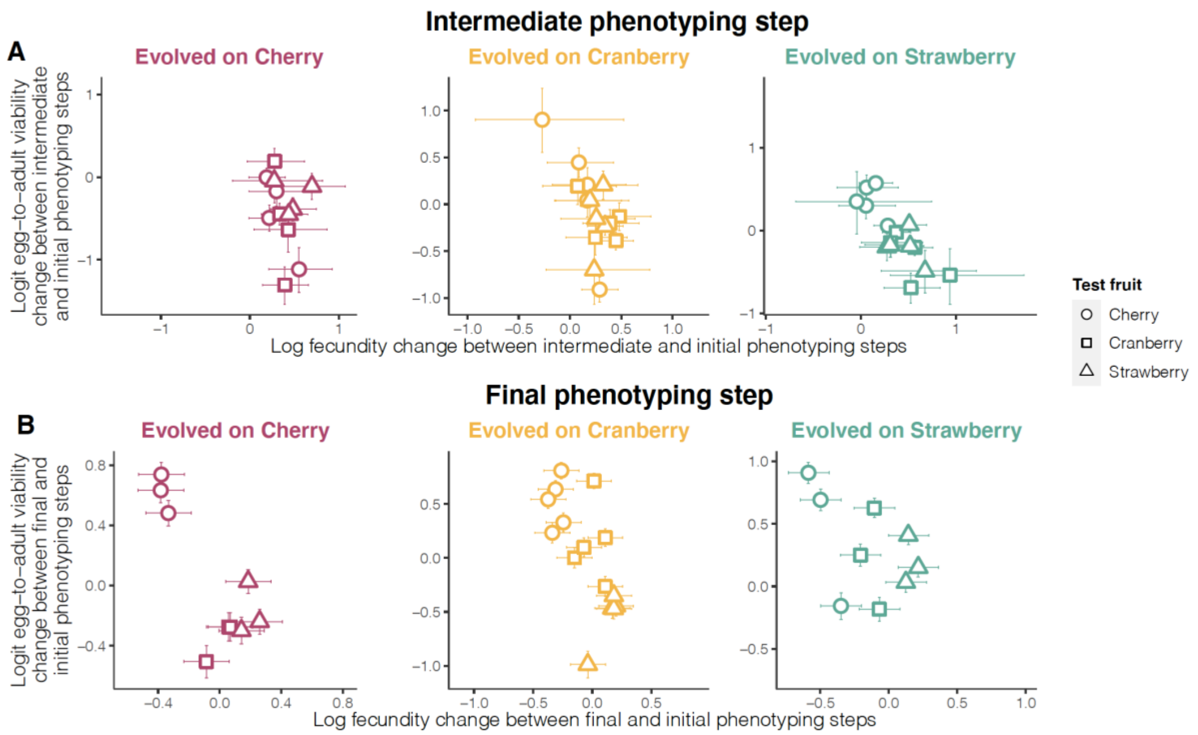


**Figure S6. Correlation between changes in fecundity and changes in egg-to-adult viability (A) between the initial and intermediate phenotyping steps and (B) between the initial and final phenotyping steps**.

### Appendix S3: Performance of the *F*-test for local adaptation (Blanquart et al., 2013) when applied to Poisson distributed data

**Motivation**

The *F*-test proposed by Blanquart et al. (2013) to test for local adaptation assumes that the fitness-related trait under study is normally and independently distributed with equal variances. We here evaluated the power and robustness of the test when applied to log-transformed non-normally distributed count data simulated under a scenario mimicking our experimental setup (i.e., similar number of populations, habitats and individuals) and with different levels of local adaptation.

**Methods**

The count data $Y_{ijk}$ for the trait (i.e., the number of eggs laid or number of emerged adults that emerged) observed in population *i* on the host plant *j* for individual *k* was simulated as follows:

$$Y_{ijk} \sim \text{Poisson}(\lambda_{ijk}) \qquad (7),$$

where $\lambda_{ijk} = \exp(\mu + s*I_{ij} + a_i + b_j + c_{ij} + \varepsilon_{ijk})$.

$\mu$ is a constant term corresponding to the overall mean counts measured in a log scale, $a_i \sim N(0,\sigma^2_a)$ is the population *i* effect, $b_j \sim N(0,\sigma^2_b)$ is the habitat *j* effect, $c_{ij} \sim N(0,\sigma^2_c)$ is the population by habitat interaction effect, and $\varepsilon_{ijk} \sim N(0,\sigma^2_\varepsilon)$ is an error term that introduces overdispersion among individuals sampled from population *i* sampled in habitat *j*. The binary auxiliary variable $I_{ij}$ indicates whether the population/habitat combination is allopatric ($I_{ij} = 0$

when $i \neq j$) or sympatric ($I_{ij}$ = 1 when $i = j$). s is the magnitude of the fitness advantage of being in sympatry (relative to allopatry).

To mimic our experimental design, we simulated I = 11 populations and J = 3 habitats and the per population/habitat combination sample size was set by default to n=30. Similarly, in our experiment, we estimated $\mu$= 3.053, $\sigma_a$= 0.165, $\sigma_b$= 0.0636; $\sigma_c$= 0.345; $\sigma_\varepsilon$ = 0.868 and s = 0.295 for the number of adults emerged during the final phenotyping. We thus considered these estimations, as default values, for the corresponding simulation parameter values. We also simulated a range of values for four parameters that were each modified one at a time (i.e., other simulation parameter being set to their default values): (i) s = 0, 0.1, 0.2, 0.3, 0.4 or 0.5 to evaluate the power of the method as a function of the magnitude of local adaptation with; (ii) $\sigma_\varepsilon$ = 0, 1, 2 or 3 to evaluate the impact of overdispersion (from absent to three times as high as the one we observed); (iii) $\mu$ = 0, 1, 2, 3, 4 or 5 to evaluate the effect of the overall mean count (from n = 1 to ca. 150 on a natural scale); and (iv) the sample size per population/habitat combination was set to 2, 10, 20 or 30 to evaluate the effect of the number of replicates per combination.

For each simulation scenario, 5,000 datasets were generated and analyzed as described in the main text. To compare the performance of the model for different parameter values, we used the R package *PRROC* (Grau et al. 2015) to compute for various p-value thresholds the (i) true positive rates (TPR) or power which corresponds to the proportion of data sets with s > 0 among the ones declared significant for local adaptation); and (ii) false positive rates (FPR) which corresponds to the proportion of data sets with s = 0 among the ones declared non-significant for local adaptation. From these estimates, standard receiver operating curves (ROC) plotting TPR against FPR could then be drawn and the area under the ROC cure (AUC) computed. Note that AUC = 1 corresponds to an optimal classifier.

**Results**

Figure S7 shows that the distribution of p-values obtained after analyzing data sets simulated with s = 0 (no local adaptation) is uniform. The Blanquart F test applied to log-transformed count data is thus well calibrated under the null hypothesis of no local adaptation, at least under the conditions of our experimental set-up.

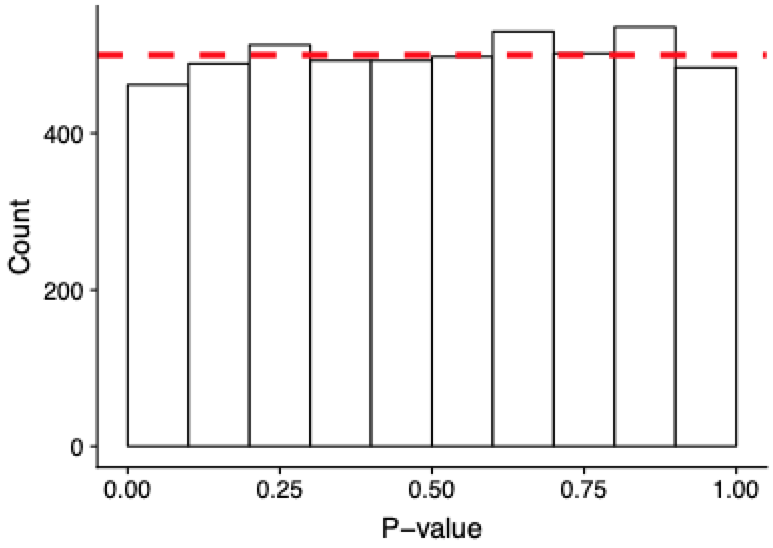


**Figure S7. Distribution of *p*-values calculated using Blanquart et al. (2013)'s F-test processed after a log transformation of data simulated following our experimental setup conditions and with a local adaptation s value equals to 0**.

The ROC curves obtained from the analysis of data sets simulated with varying magnitude of local adaptation (from s = 0.1 to s = 0.5) are plotted on Figure S8. As expected, the performance of the model improved with s, the ROC-AUC being above > 0.99 for s ≥ 0.3, the latter value being similar to the one we estimated on real data in our experiment.

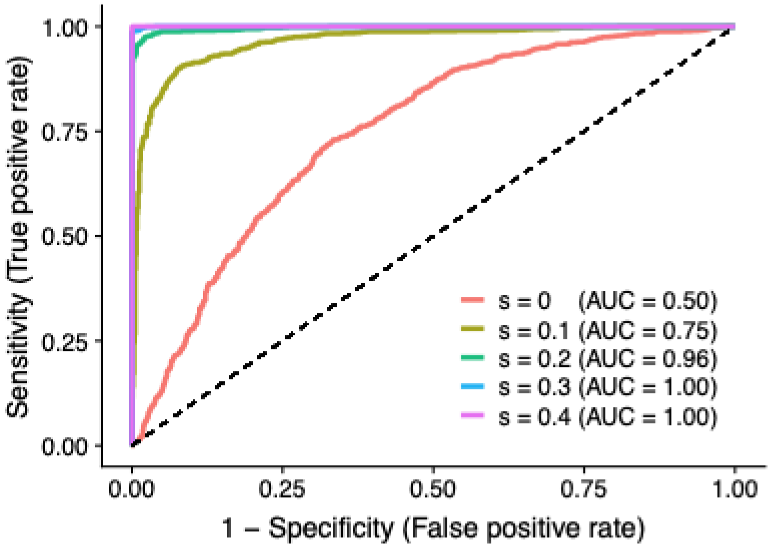


**Figure S8. Evaluation of the performance of the method for varying magnitude of local adaptation (measured by s).**
Count data sets were simulated with five different values of s ranging from s = 0.1 to 0.5 (other simulation parameters being set to their default value). In each case, 5,000 data sets were then analyzed with the *F*-test by Blanquart et al. (2013) after log-transformation to estimate TPR and FPR (averaged over all the data sets). The corresponding ROC curves are plotted and ROC-AUC are given in parentheses in the figure legend.

Figure S9 gives the ROC curves obtained from the analyses of simulated data sets when varying i) the population by habitat sample size (n = 2 to n = 30); ii) overall mean count (from $\mu = 0$ to $\mu = 5$ in log-scale); and iii) overdispersion (from $\sigma_\varepsilon = 0$ to $\sigma_\varepsilon = 3$). The approach was mainly found sensitive to a smaller number of individuals (n < 20) and to higher overdispersion ($\sigma_\varepsilon > 2$) than the ones corresponding to our experiment. Interestingly, the overall average mean count had only a minor effect on the performance of the method.

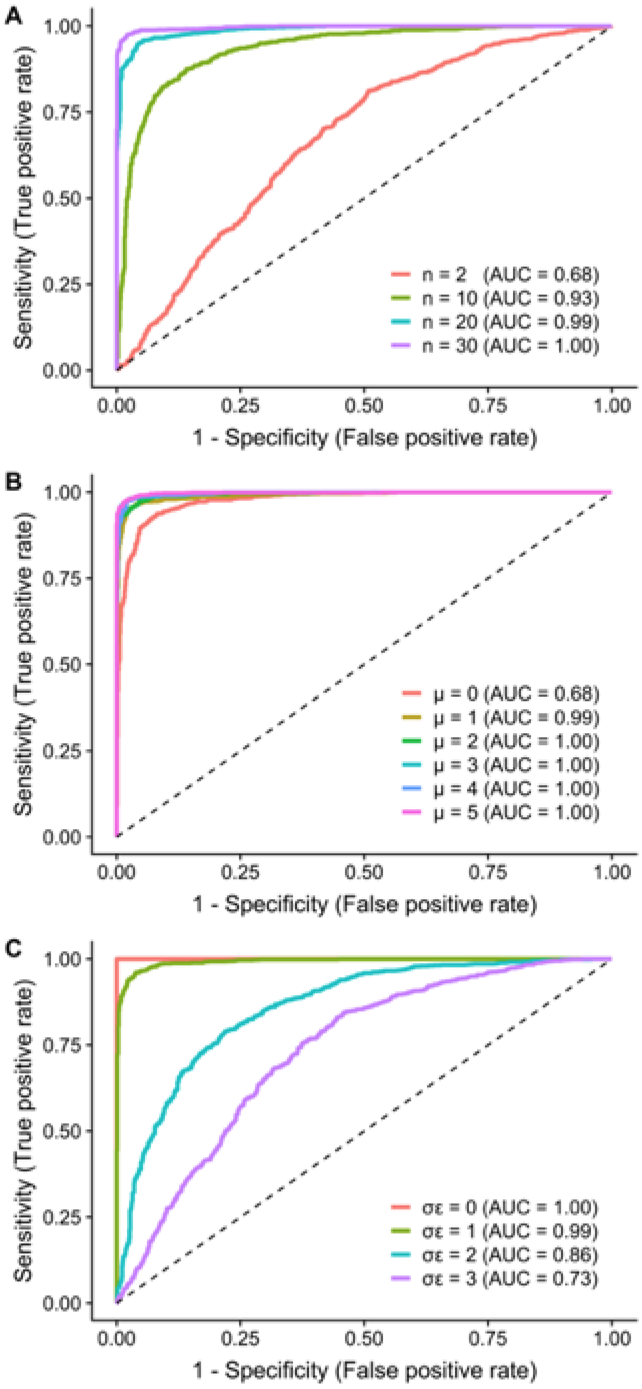


**Figure S9. Evaluation of the performance of the method when varying (A) the population by habitat sample size (measured by n), (B) the overall mean count (measured by μ) and (C) the overdispersion of data (measured by $\sigma_\varepsilon$).**

Count data sets were simulated with four different values of n ranging from n = 2 to 30 (other simulation parameters being set to their default value), six different values of μ ranging from n = 0 to 5 (other simulation parameters being set to their default value), and four different values of $\sigma_\varepsilon$ ranging from n = 0 to 3 (other simulation parameters being set to their default value). In each case, 5,000 data sets were then analyzed with the *F*-test of Blanquart et al. (2013) after log-transformation to estimate TPR and FPR (averaged over all the data sets). The corresponding ROC curves are plotted and ROC-AUC are given in parentheses in the figure legend.

#### General conclusion

From the power analysis results given above, we conclude that the F-test proposed by Blanquart et al. (2013) to detect local adaptation can be applied to log-transformed count data provided that the population by habitat sample size is high enough (> 20) and overdispersion of the data remained limited, as previously discussed in O’Hara et Kotze (2010).

### Appendix S4: Effect of the temporary pooling of replicate populations on the negative association between fitness changes in selective and alternative environments

It is difficult to know whether the temporary pooling of replicate populations is responsible for the similarity of fitness changes among replicate populations evolving on the same fruit medium. To investigate this potential confounding effect, we identified and tested two potential scenarios for the effect of pooling on the fitness changes observed in populations evolved on cranberry or cherry.

In the first scenario, the different populations contributed equally to the pool. For each fruit medium, the fitness of the population after pooling is equal to the mean of the fitnesses of the different populations before pooling. Figure S10 shows that under this scenario pooling does not result in negative correlations between fitness changes in selective and alternative environments for populations evolved on cranberry or cherry.

In the second scenario, the population with the highest fitness among populations evolving on the same fruit medium contributed disproportionately more to the pool. In other words, for each fruit medium, the fitness of the population after pooling is equal to the fitness of the population with highest mean fitness. Figure S11 shows that under this scenario pooling does not result in negative correlations between fitness changes in selective and alternative environments for populations evolved on cranberry or on cherry media. Hence, it seems unlikely that the temporary pooling of replicate populations generated the pattern observed during the final phenotyping step.

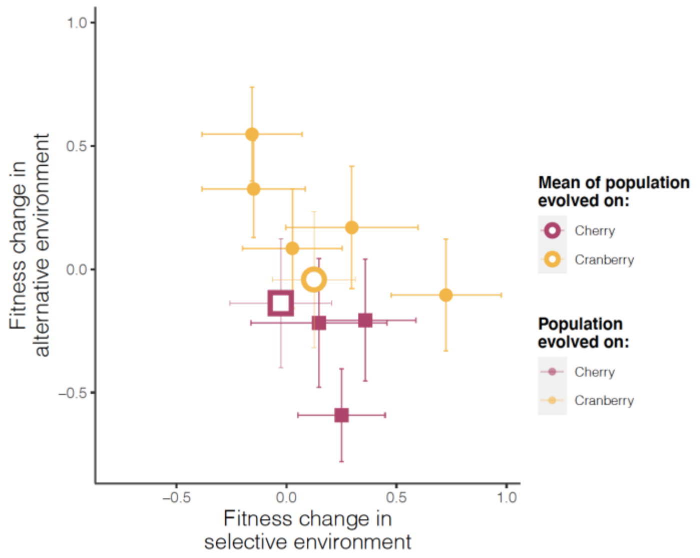


**Figure S10: First scenario for the effect of pooling on the emergence of a negative association between fitness changes in selective and alternative environments (fitness of the population after pooling is equal to the mean of the fitness of the different populations before pooling)**. Open circles represent the average fitness change before pooling of the different replicate populations that evolved on the same selection fruit. Closed circles represent the fitness change between final and initial phenotyping steps measured on selective and alternative fruit media for cherry and cranberry. Error bars represent 95% confidence intervals.

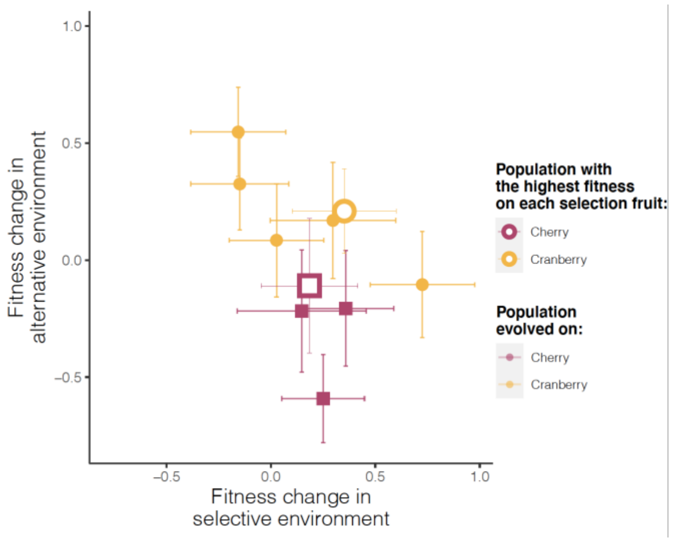


**Figure S11: Second scenario for the effect of pooling on the emergence of negative genetic correlations (fitness of the population after pooling is equal to fitness of the population with highest fitness on each selection fruit).** Open circles represent the average fitness change of the population with the highest fitness on each selection fruit during the intermediate phenotyping. Closed circles represent the fitness change between final and initial phenotyping steps measured on selective and alternative fruit media for cherry and cranberry. Error bars represent 95% confidence intervals.
